# Spatial dynamics of the tumor microenvironment linked to emerging resistance in *EGFR*-mutated lung cancer

**DOI:** 10.1101/2025.02.10.637543

**Authors:** Satoshi Nakamura, Daisuke Shibahara, Kentaro Tanaka, Yasuyuki Kishikawa, Mikiko Hashisako, Keita Nakatomi, Noriaki Nakagaki, Mikihiro Kohno, Koichi Azuma, Ritsu Ibusuki, Kohei Otsubo, Yasuto Yoneshima, Eiji Iwama, Yoshinao Oda, Isamu Okamoto

## Abstract

Epidermal growth factor receptor (EGFR) tyrosine kinase inhibitors (TKIs) provide therapeutic benefit in *EGFR* mutation–positive non–small cell lung cancer, but some individuals develop early resistance. We performed spatial transcriptomics analysis of pre- and posttreatment tumor samples from the same patients to explore the underlying mechanisms of such early resistance. The proportion and activation of fibroblasts increased in association with the development of early resistance, whereas a distinct tumor cell cluster showed activation of tumor necrosis factor–α signaling via the NF-κB pathway even before treatment. Also in the early resistance sample, specific tumor cell clusters interacted with immune and stromal cells. Immature tertiary lymphoid structures (TLSs) were enriched in the early resistance sample, whereas mature TLSs were observed in the long-term response sample. These findings implicate tumor heterogeneity and an inflammatory tumor microenvironment in early EGFR-TKI resistance, providing insight into potential therapeutic strategies to improve treatment outcomes.

## Introduction

Epidermal growth factor receptor (EGFR) tyrosine kinase inhibitors (TKIs) show durable therapeutic efficacy in most individuals with non–small cell lung cancer (NSCLC) harboring activating mutations of the *EGFR* gene^1^. Nevertheless, resistance to EGFR-TKIs inevitably emerges in such patients after treatment initiation^2,3^. Whereas several mechanisms of resistance to EGFR-TKIs have been identified, the underlying cause of resistance remains unknown in 40% to 50% of cases^4–6^. Furthermore, some patients develop early resistance during EGFR-TKI treatment, whereas others achieve a long-term response, with the reasons for these divergent outcomes also remaining unclear. There is therefore a pressing need to elucidate the precise mechanisms of EGFR-TKI resistance not only for the development of effective therapies to overcome such resistance but also to extend the duration of the response to EGFR-TKI treatment. Increasing evidence has suggested that the tumor microenvironment (TME) plays a pivotal role in shaping the response to EGFR-TKIs^7–12^. However, spatial analyses of the TME based on paired pre- and posttreatment tumor specimens from the same patients with *EGFR*-mutated NSCLC have been limited^13^.

Spatial transcriptomics analysis has emerged as a powerful approach to characterization of the TME^13–18^. As an array-based method, Visium HD (10× Genomics) allows both comprehensive mRNA capture, without limiting the analysis to specific target genes, as well as analysis of an entire specimen section. We have now performed a detailed analysis with Visium HD of the TME in paired pre- and posttreatment tumor specimens from patients with *EGFR*-mutated NSCLC. Furthermore, we converted Visium HD spot data into single-cell data with the use of StarDist, a deep learning–based method for detection and segmentation of nuclei and cells. By integrating the spatial coordinates derived from StarDist into the single-cell data, we were able to conduct spatial transcriptomics analysis at the single-cell level while preserving the actual spatial relations between cells. Our findings provide insight into the spatial dynamics of the TME, highlighting a contribution of an inflammatory TME and tumor heterogeneity to the early onset of EGFR-TKI resistance in *EGFR*-mutated NSCLC.

## Results

### Integrative analysis of spatial transcriptomics and nuclear segmentation

To investigate differences in the duration of response to osimertinib, a third-generation EGFR-TKI, we collected and analyzed paired pre- and posttreatment tumor specimens from four individuals with *EGFR*-mutated NSCLC (Fig. 1A and fig S1). These individuals were categorized as one with early resistance (ER), one with a standard response (SR), and two with a long-term response (LR) on the basis of progression-free survival (PFS)^2^, and their clinical characteristics are provided in table S1. All samples were processed with the Visium HD platform, which provides a spatial resolution of 2 µm per spot. To reconstruct single cells from the data, we applied StarDist^19^, a deep learning–based tool that allows the detection and segmentation of nuclei from hematoxylin-eosin (H&E)–stained images generated alongside the spatial transcriptomics data (Fig. 1B and fig. S2). With the use of StarDist, we aggregated 2-µm spots that overlapped with the polygonal coordinates corresponding to the detected nuclei, thereby reconstructing spatial transcriptomics data mapped to individual cell positions (Fig. 1C). In addition, we incorporated the centroid coordinates of each polygon as new spot coordinates into the data structure to ensure compatibility with existing tools for spatial transcriptomics analysis (fig. S3, A to C).

**Fig. 1.**
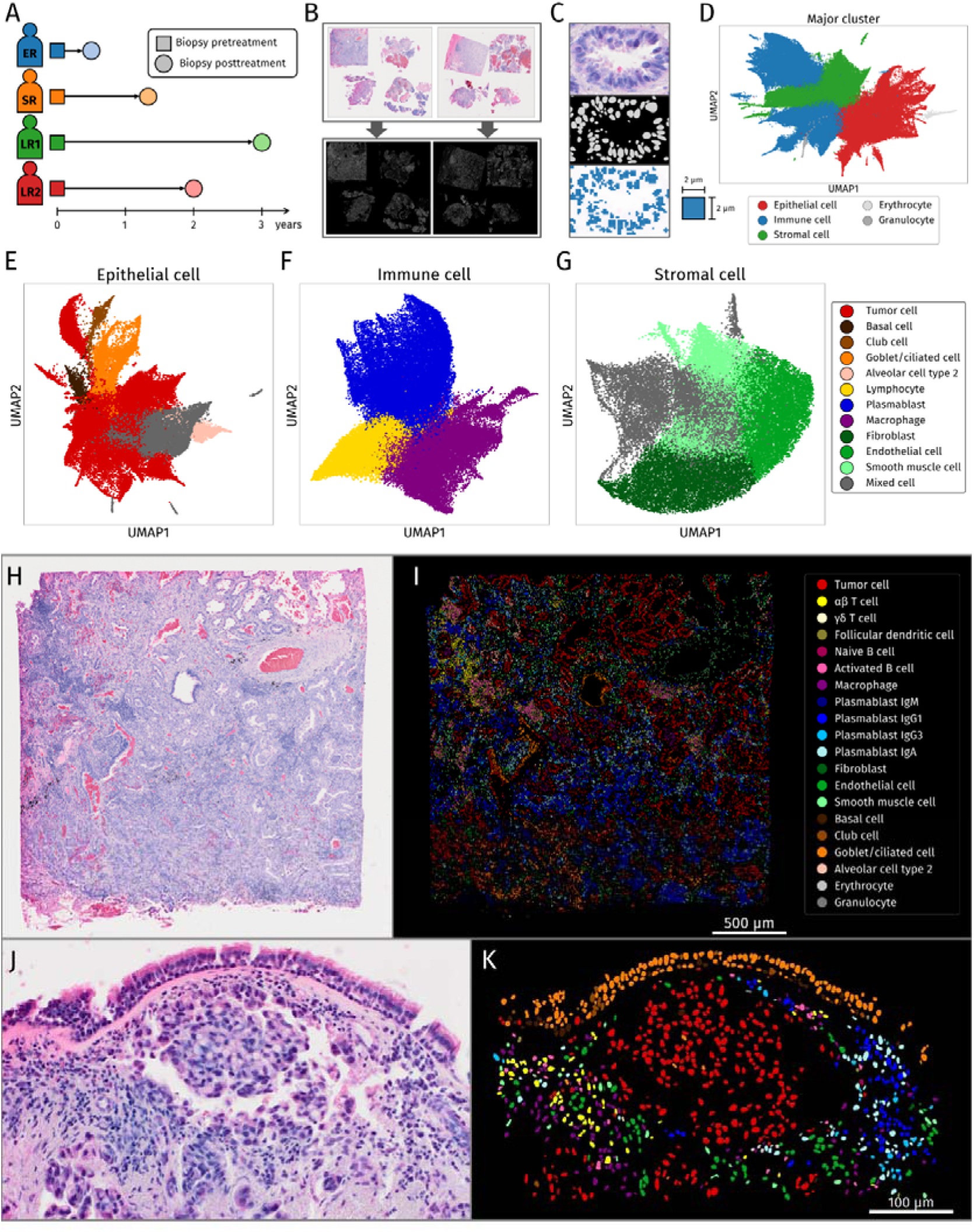
Analysis overview, clustering, and visualization of cell type and spatial location. (**A**) Biopsy interval for pre– and post–osimertinib treatment and patient classification. (**B**) Detection and segmentation of cell nuclei with StarDist on high-resolution tissue images loaded with the Visium HD protocol. (**C**) Visualization of detection and segmentation by StarDist and 2-µm spot extraction for identified cell nuclei positions. (**D**) UMAP of annotated cell types from single-cell RNA-seq analysis based on cell nuclei identified by StarDist. (**E** to **G**) UMAP with epithelial cells (E), immune cells (F), and stromal cells (G) reclustered and labeled for cell type. (**H** and **I**) H&E-stained pathological image of an area of the ER_pre sample (H) and image of the same area with cell type identification reflected in polygonal coordinate data obtained by StarDist (I). (**J** and **K**) H&E-stained pathological image for a magnified region of the LR2_pre sample (J) and image of the same region with cell type identification reflected in polygonal coordinate data obtained by StarDist (K). Normal epithelial cells and tumor cells were distinguished, and surrounding mesenchymal cells and immune cells were identified with high accuracy.

A total of 814,636,020 reads were obtained from Slide 1 (ER, SR), and 730,477,318 reads were obtained from Slide 2 (LR1, LR2). StarDist identified 141,324 nuclei in Slide 1 and 99,658 nuclei in Slide 2. Samples were partitioned on the basis of coordinates for their location on the image, and the sample names were linked to the data (fig. S3D). The integrated data for the eight samples were then processed for quality control, filtering, normalization, dimensionality reduction, batch correction, nearest-neighbor graph building, and Leiden clustering, and they were visualized by uniform manifold approximation and projection (UMAP). Cells were broadly categorized into three main lineages—epithelial cells, stromal cells, and immune cells—together with a small number of blood cells (Fig. 1D and Fig. S3E). The epithelial cell population was further divided into tumor cells, basal cells, club cells, alveolar cells type 2, and goblet/ciliated cells (Fig. 1E and Fig. S4A). Immune and stromal cells were clustered into lymphocytes, plasmablasts, macrophages, fibroblasts, endothelial cells, and smooth muscle cells (Fig. 1, F and G, and fig. S4, B and C). In addition, lymphocytes were subdivided by type, plasmablasts were classified by the immunoglobin (Ig) they produced, and macrophages were distinguished from mixed cells (Fig. S4, D to I). The results of these clustering analyses were integrated with the polygonal coordinate data obtained from StarDist, thereby allowing visualization of distinct cell populations on spatial coordinates (Fig. 1, H to K, and fig. S5 and S6).

### Changes in the TME between pre- and posttreatment samples

To investigate changes in tumor cell characteristics between before and after treatment, we performed differential gene expression (DGE) analysis following pseudo–bulk processing, comparing tumor cells from pre- and posttreatment samples (Fig. 2A). Genes with significantly higher expression in the pretreatment group showed a tendency to be associated with a favorable prognosis for NSCLC patients in the Kaplan-Meier Plotter database (fig. S7, A to D), whereas those with higher expression in the posttreatment group tended to be associated with a poorer prognosis (fig. S7. E to H). Gene set enrichment analysis (GSEA) revealed significant enrichment of gene sets associated with the cell cycle and cell proliferation in posttreatment tumor cells (Fig. 2, B and C, and fig. S7. I and J). These results suggested that tumor cells had an enhanced proliferative activity after the development of EGFR-TKI resistance.

**Fig. 2.**
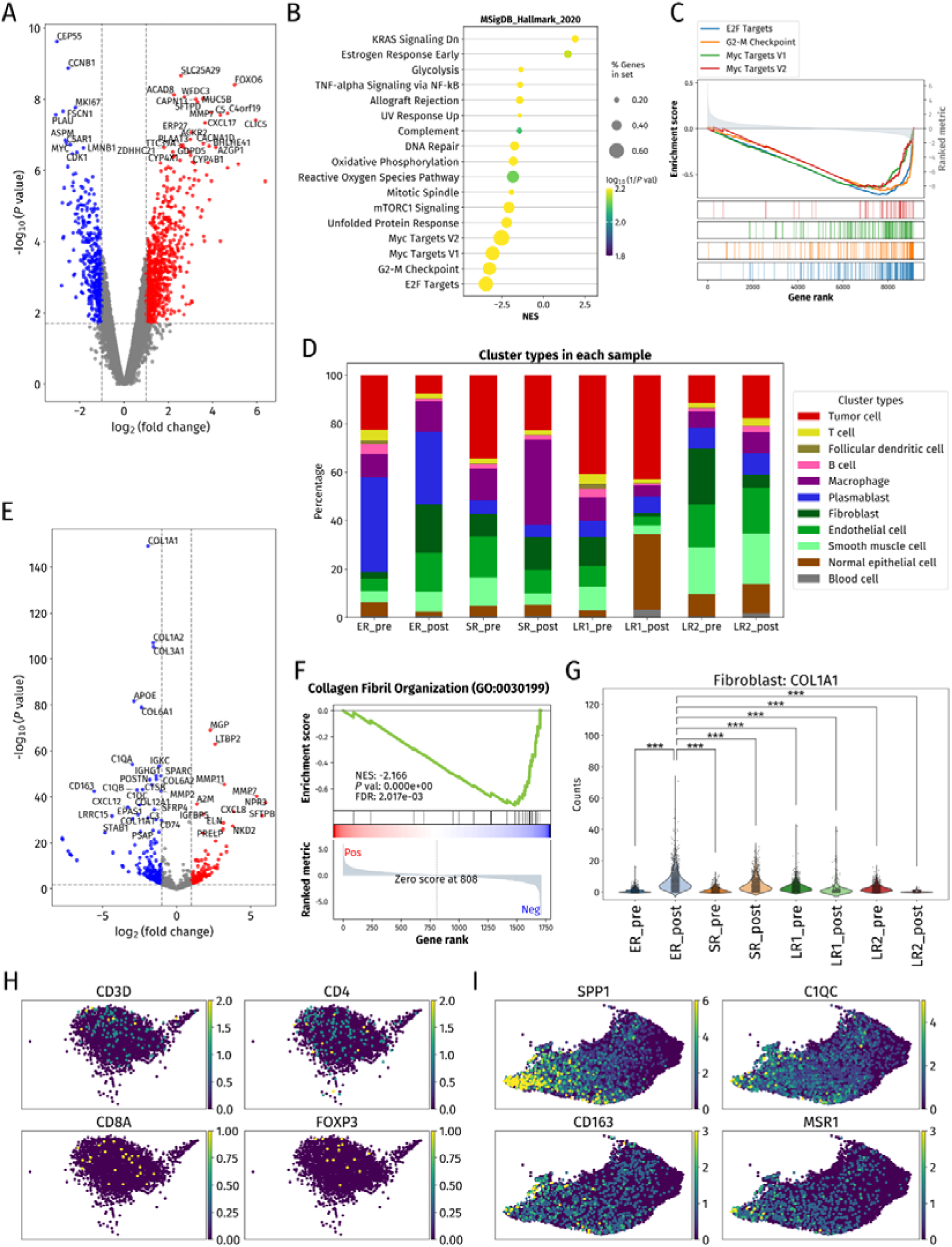
Comparative analysis of samples obtained before and after treatment. (**A**) Volcano plot comparing tumor cell clusters from all samples dividing into two groups corresponding to before and after treatment (All_pre vs All_post). The red and blue dots indicate significantly upregulated genes of All_pre and All_post, respectively. The vertical dashed lines represent the fold change of −2 and 2, and the horizontal dashed line represents the *P* value of 0.05. (**B**) Dot plot for GSEA with Hallmark gene sets based on DGE analysis of tumor cells for All_pre vs All_post. (**C**) GSEA plots for gene sets with high negative values of the normalized enrichment score (NES) for tumor cells in All_pre vs All_post. (**D**) Stacked bar chart showing the proportion of cell types within each sample. (**E**) Volcano plot for comparison of fibroblasts extracted from ER_pre and ER_post samples. (**F**) GSEA plot of the only significant gene set among Hallmark gene sets for fibroblasts in ER_pre vs ER_post. FDR, false discovery rate. (**G**) Violin plot and strip plot showing raw counts for the *COL1A1* gene in fibroblasts of each sample. *\*\*\*P* < 0.0005 (one-sided Mann-Whitney U test). (**H**) Expression levels of *CD3D*, *CD4*, *CD8A*, and *FOXP3* related to T cell subtypes shown on the UMAP plot of T cell parts extracted from lymphocyte clusters. (**I**) Expression levels of genes related to tumor-associated macrophages (*SPP1*, *C1QC*) and M2 macrophages (*CD163*, *MSR1*) on the UMAP plot of macrophages.

We next compared the proportions of stromal and immune cell clusters between pre- and posttreatment samples. The proportion of stromal cells, including fibroblasts and endothelial cells, was increased in the ER_post sample compared with the ER_pre sample (Fig. 2D). DGE analysis of fibroblasts from the ER samples revealed increased expression of collagen-related genes after treatment. (Fig. 2E). GSEA further revealed that a gene set related to extracellular matrix productionwas significantly enriched in fibroblasts of the ER_post sample (Fig. 2F). In addition, collagen gene expression in fibroblasts was significantly increased for the ER_post sample compared with all other samples (Fig. 2G and fig. S7K). These results suggested that the activity of fibroblasts was increased after EGFR-TKI treatment, contributing to the changes in the TME associated with early resistance.

Analysis of the T cell cluster revealed that CD4-expressing T cells were present in slightly higher numbers than CD8-expressing or regulatory T cells, although low expression levels of *CD4*, *CD8A*, and *FOXP3* impeded the classification of these cells (Fig. 2H and fig. S7L), possibly as a result of the filtering effect of the analytic methods applied. For the macrophage cluster, tumor-associated macrophage markers (*SPP1*, *C1QC*) were highly expressed in accordance with the distribution of M2-like markers (*CD163*, *MSR1*)^20–22^, suggestive of an immune-suppressive phenotype that was consistently observed across all samples (Fig. 2I and fig. S7M).

### Spatial distribution of tumor cell heterogeneity associated with early resistance or a longer-term response

To explore factors that might contribute to early resistance, we categorized tumor cells from all pretreatment samples into two groups: ER_pre and SLR_pre (the latter comprising SR_pre, LR1_pre, and LR2_pre) for DGE analysis (Fig. 3A). GSEA revealed significant enrichment of gene sets related to the inflammatory response, tumor necrosis factor–α (TNF-α) signaling via the nuclear factor–κB (NF-κB) pathway, the interleukin-6 (IL-6) pathway, and chemokine signaling pathways in ER_pre tumor cells (Fig. 3, B and C, and fig. S8, A and B), indicating that the TME of ER_pre was characterized by active inflammation.

**Fig. 3.**
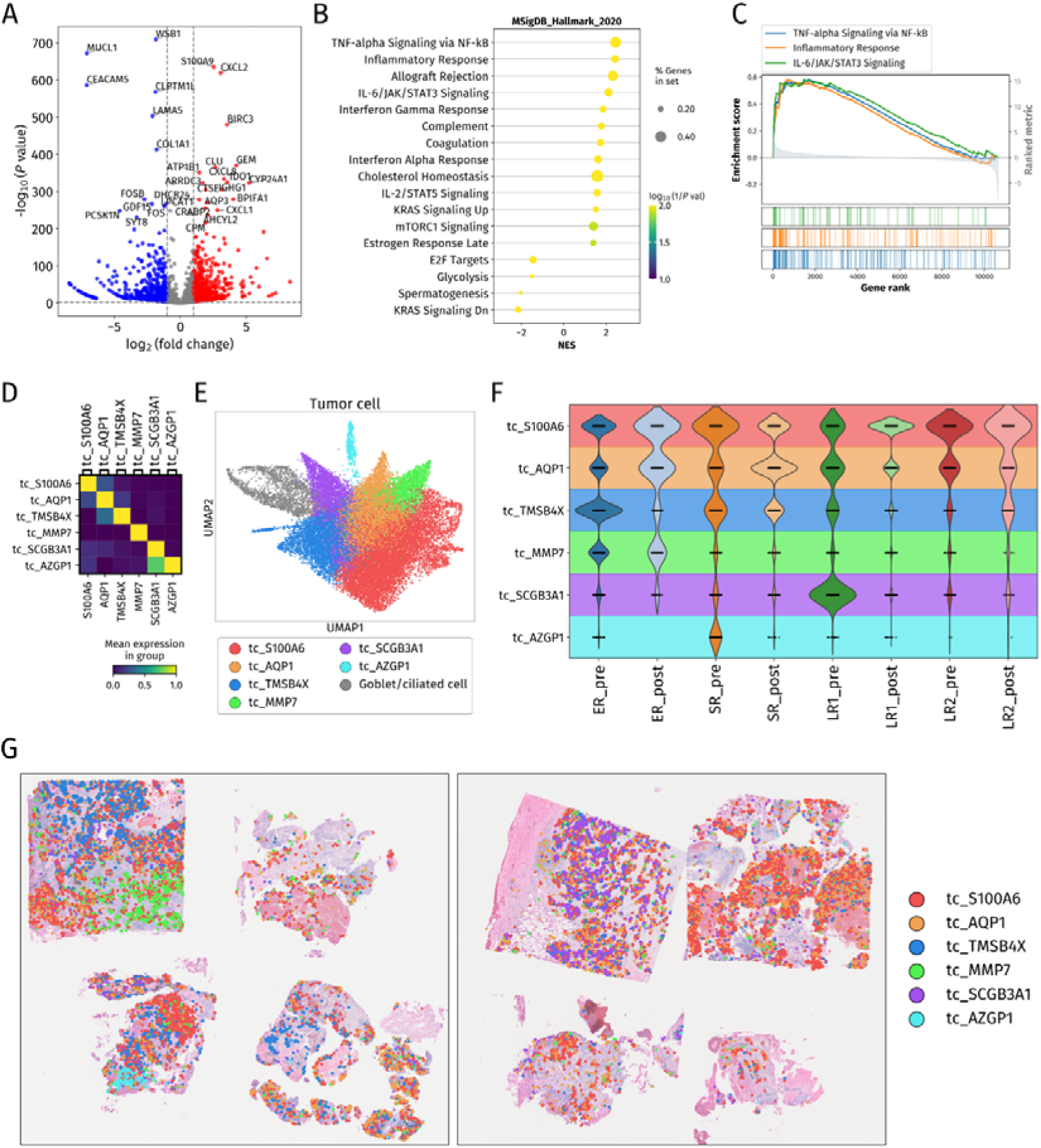
Analysis exploring mechanisms underlying tumor heterogeneity and early resistance. (**A**) Volcano plot for DGE analysis results obtained by extracting tumor cell clusters from pretreatment samples and dividing them into two groups: ER_pre and SLR_pre. (**B**) Dot plot for GSEA with Hallmark gene sets based on DGE analysis of ER_pre vs SLR_pre. (**C**) GSEA plots for gene sets with high positive values of NES for ER_pre vs SLR_pre. (**D**) Matrix plot showing clustering of tumor cells based on differentially expressed cancer-related genes. (**E**) UMAP plot with tumor cells reclustered and labeled for differentially expressed genes. (**F**) Violin plot for the distribution of tumor cell clusters in each sample (**G**) Image of tumor cell cluster positions plotted on H&E-stained tissue images for Slide 1 and Slide 2.

To better understand the factors that contribute to the differences between samples, we performed a comparative analysis of tumor cell clusters. We classified and labeled each tumor cell cluster on the basis of DGE for cancer-related genes^23–28^ as discriminatory markers (Fig. 3, D and E, and fig. S8. C and D). Application of inferCNV for analysis of copy number variation (CNV) revealed a tendency for consistent CNV alterations in each tumor cell cluster in comparison with other normal epithelial cell clusters as a reference group (fig. S8E). To quantify their distribution, we visualized the tumor cell clusters across samples and mapped them spatially for each sample (Fig. 3, F and G). The tc_S100A6 and tc_AQP1 clusters were present in all pre-and posttreatment samples, accounting for a high proportion of tumor cells in most samples. The tc_S100A6 cluster manifested significantly higher expression of *CCND1* and *ERBB2* compared with the other clusters (fig. S8, F anf G), whereas the tc_AQP1 and tc_TMSB4X clusters showed increased expression of *MET* and *CD74*, respectively (fig. S8, H and I). GSEA based on DGE analysis of the tc_S100A6 cluster revealed significant enrichment of gene sets related to cell proliferation, including E2F targets and the G2-M checkpoint, compared with the other clusters (Fig. 4, A and B). In contrast, gene sets related to the inflammatory response, epithelial-mesenchymal transition (EMT), and KRAS signaling were not enriched (Fig. 4C). Similar DGE analysis and GSEA for the tc_AQP1 cluster revealed upregulation of gene sets associated with mechanistic target of rapamycin complex 1 (mTORC1) and IL-6–JAK–STAT3 signaling, both of which have been implicated in EGFR-TKI resistance^4,29–31^, in addition to those related to EMT and the inflammatory response (Fig. 4, D to F). We next analyzed distinct clusters that showed differences between samples.

**Fig. 4.**
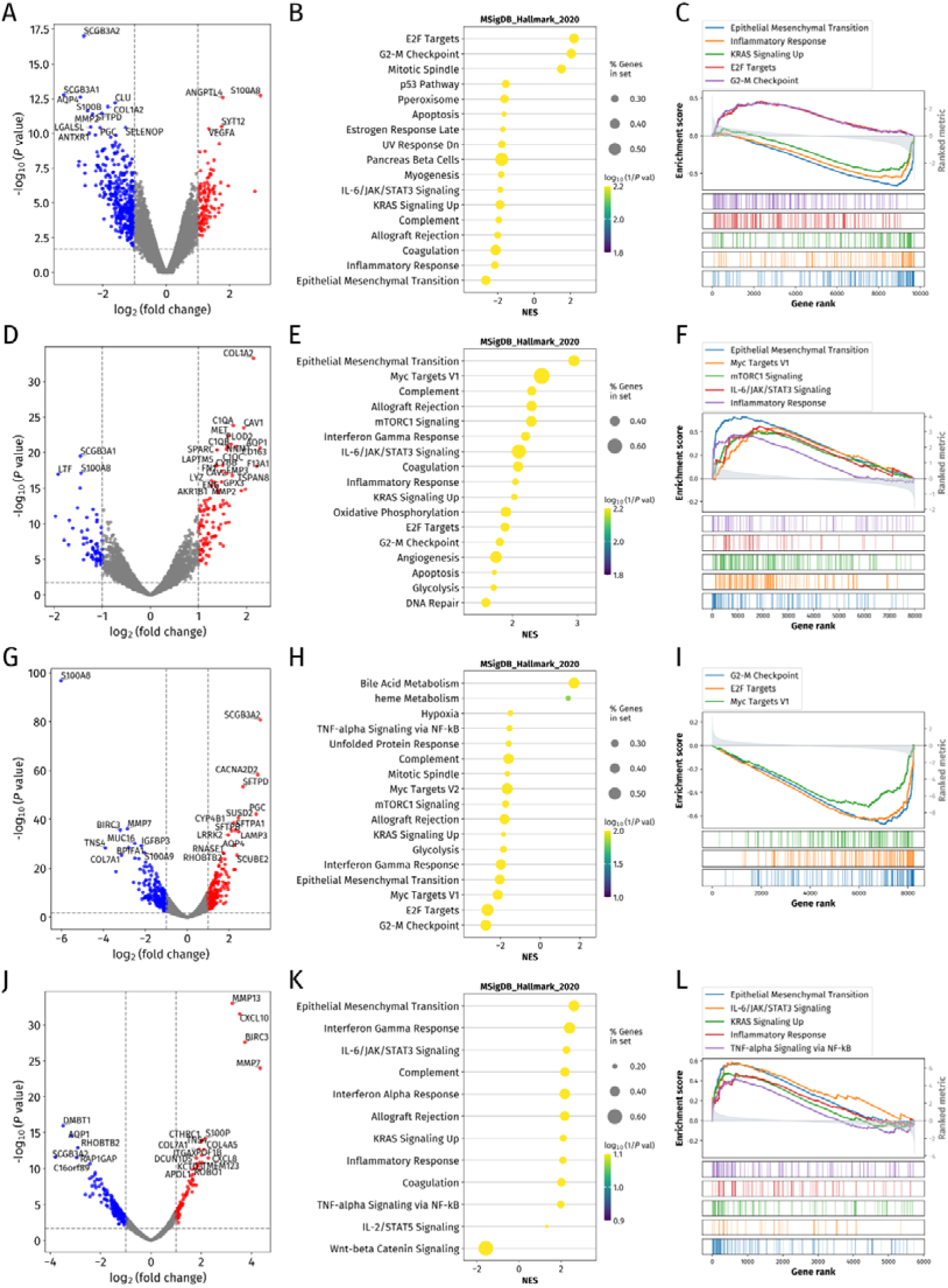
Analysis of individual tumor cell clusters. (**A**) Volcano plot of DGE analysis results for the tc_S100A6 tumor cell cluster versus the corresponding other tumor cell clusters (tc_rest). (**B**) Dot plot for GSEA with Hallmark gene sets based on DGE analysis of tc_S100A6 vs tc_rest. (**C**) GSEA plots for gene sets with high absolute values of NES for tc_S100A6 vs tc_rest. (**D**) Volcano plot of DGE analysis results for tc_AQP1 vs tc_rest. (**E**) Dot plot for GSEA with Hallmark gene sets based on DGE analysis of tc_AQP1 vs tc_rest. (**F**) GSEA plots for gene sets with high positive values of NES for tc_AQP1 vs tc_rest. (**G**) Volcano plot of DGE analysis results for tc_TMSB4X vs tc_rest. (**H**) Dot plot for GSEA with Hallmark gene sets based on DGE analysis of tc_TMSB4X vs tc_rest. (**I**) GSEA plots for gene sets with high negative values of NES for tc_TMSB4X vs tc_rest. (**J**) Volcano plot of DGE analysis results for tc_MMP7 vs tc_rest. (**K**) Dot plot for GSEA with Hallmark gene sets based on DGE analysis of tc_MMP7 vs tc_rest. (**L**) GSEA plots for gene sets with high positive values of NES for tc_MMP7 vs tc_rest.

The tc_TMSB4X cluster was most pronounced in the ER_pre sample (Fig. 3F), suggestive of a potential role in early resistance. DGE analysis and GSEA for this cluster revealed significant enrichment of gene sets associated with metabolic pathways, whereas those related to cell proliferation showed relatively low expression (Fig. 4, G to I). The tc_MMP7 cluster was prominent in both pre- and posttreatment ER samples but was essentially absent from the other samples (Fig. 3F), also suggestive of a potential role in early resistance. DGE analysis and GSEA for this cluster showed significant enrichment of pathways related to EMT, the inflammatory response, IL-6–JAK–STAT3 signaling, KRAS signaling, and TNF-α signaling via NF-κB (Fig. 4, J to L). The tc_SCGB3A1 cluster was most abundant in LR1_pre (Fig. 3F), possibly indicating a contribution to long-term response. DGE analysis and GSEA for this cluster revealed downregulation of gene sets related to E2F targets and the G2-M checkpoint, suggestive of a relatively slow cell cycle (fig. S9, A to C). The tc_AZGP1 cluster was characteristic of the SR_pre sample (Fig. 3F), but no significant enrichment of gene sets was observed for this cluster (fig. S9, D to F), possibly as a result of its extremely low cell number.

To investigate differences in the TME across tumor cell clusters, we quantified the spatial distribution of surrounding cell types within the ER_pre sample. For each tumor cell cluster, we calculated the proportion of cell types within concentric annular regions extending up to 1000 μm from each tumor cell (fig. S9G). This analysis revealed that the tc_S100A6 and tc_AQP1 clusters were diffusely distributed, whereas the tc_TMSB4X and tc_MMP7 clusters were each localized in close proximity, a pattern also observed in the spatial mapping of ER_pre (Fig. 3G). Of note, the tc_MMP7 cluster showed the highest proportion of immune cells within its surrounding microenvironment, consistent with the enrichment of genes associated with the inflammatory response in this cluster revealed by GSEA (Fig. 4L). Together, these findings suggested that this cluster is characterized by an inflammatory state.

### Maturity of tertiary lymphoid structures in the TME

To examine the spatial relations of each cluster within the TME, we performed a neighborhood enrichment analysis based on the spatial map coordinates annotated with cell types. This analysis revealed a close spatial relation between T cells, B cells, and follicular dendritic cells (FDCs), particularly in the ER_pre and LR1_pre samples (Fig. 5A). Visualization of these immune cell populations on the spatial map revealed that they were distributed across several regions of the ER_pre sample, whereas only one region was observed in the LR1_pre sample (fig. S10, A and B). In the ER_pre sample, these regions were characterized by a structure in which B cells and FDCs were intermixed and sparsely surrounded by T cells (Fig. 5B and fig. S10C). In contrast, in the LR1_pre sample, FDCs were positioned around B cells, with T cells surrounding the outer perimeter (Fig. 5C and fig. S10D). Moreover, *CXCL13* expression was increased within these structures (Fig. 5, B and C), indicating that they corresponded to tertiary lymphoid structures (TLSs). On the basis of previous observations^32^, we identified the poorly organized regions in the ER_pre sample as immature TLSs and the well-structured region in the LR1_pre sample as a mature TLS.

**Fig. 5.**
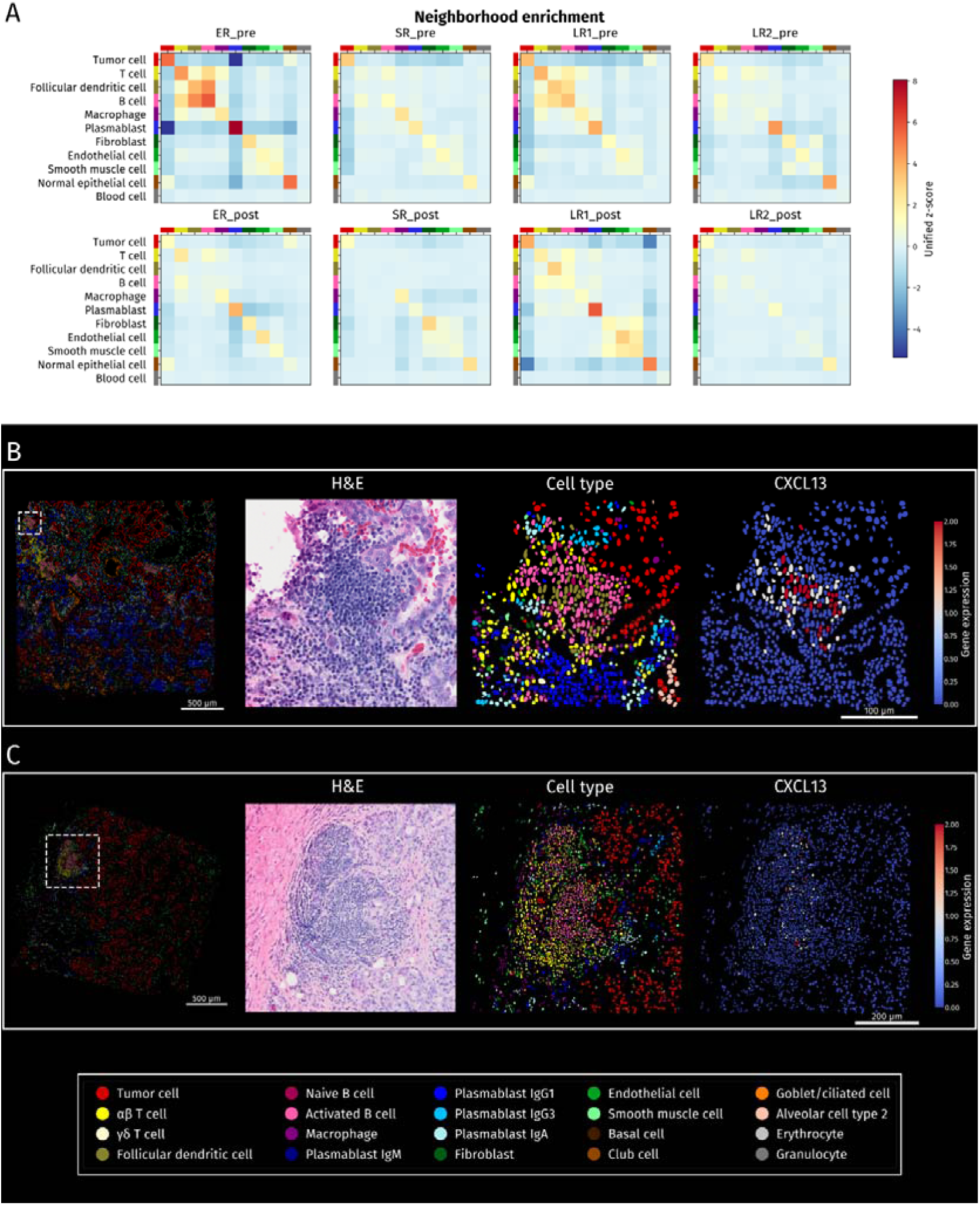
Spatial neighborhood enrichment analysis and TLS identification. (**A**) Heat maps for the results of spatial neighborhood enrichment analysis for each sample. A total of 5000 surrounding cells was counted for each cell, and the proportion of each cell type among these cells was expressed as a z-score normalized relative to all samples. (**B** and **C**) Enlarged images of a TLS in the ER_pre (B) and LR1_pre (C) samples. The area enclosed by the white dashed line is shown in the images of H&E staining, cell types, and expression of *CXCL13*.

### Enrichment of MIF- and PDGF-mediated cell-cell interactions in the TME associated with early resistance

To further elucidate the characteristics of the TME, we analyzed intercellular interactions with the use of COMMOT. This Python library calculates cell-cell interactions on the basis of ligand-receptor competition and spatial proximity. We extracted ligand-receptor gene pairs with sufficient expression levels from the CellChat database of known ligand-receptor interactions (table S2). Pathway comparisons across samples revealed increased expression of genes associated with the macrophage migration inhibitory factor (MIF) pathway in ER and SR samples (Fig. 6, A and B).

**Fig. 6.**
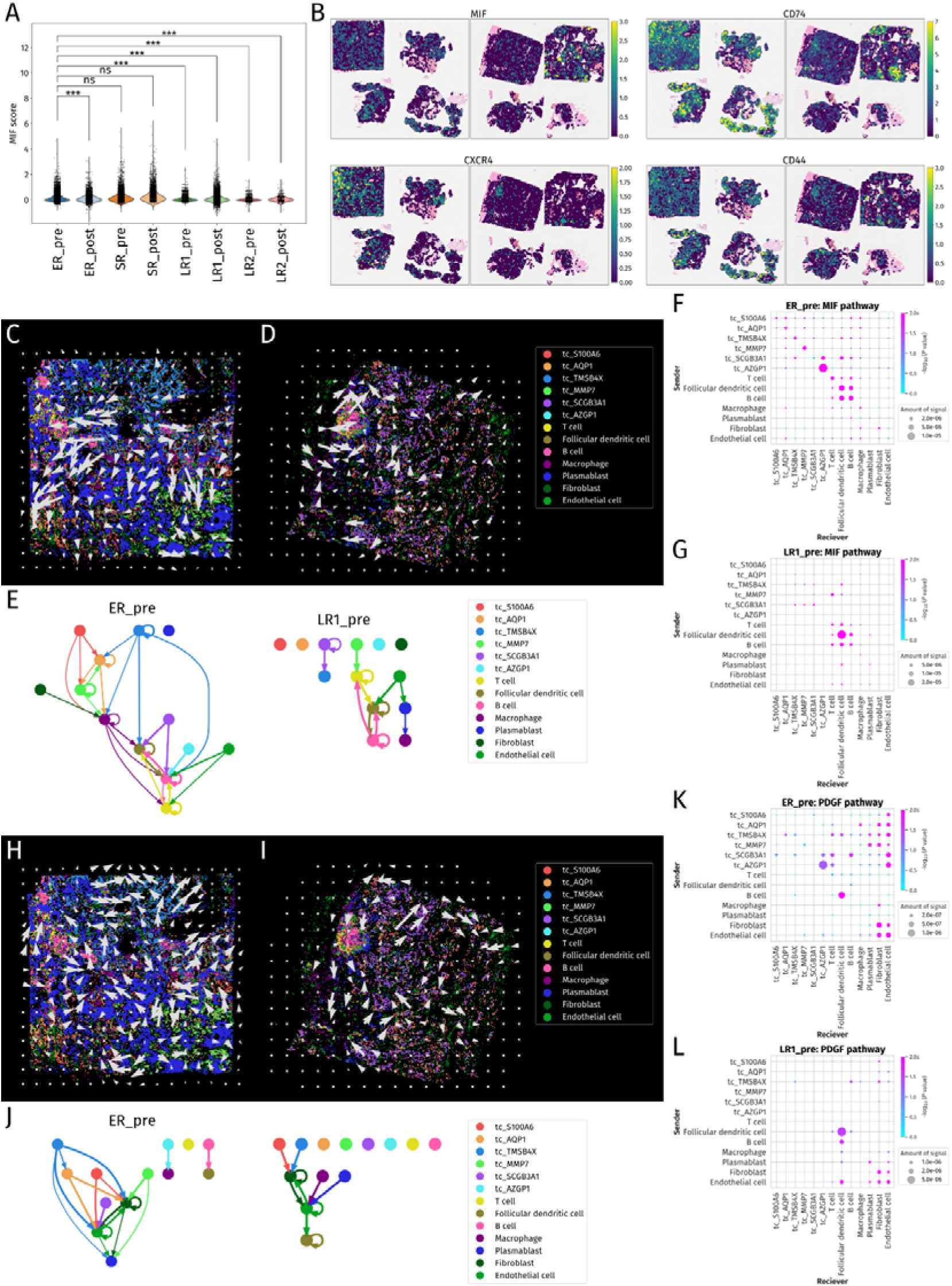
Cell-cell interaction analysis with COMMOT. (**A**) Violin plot and strip plot of MIF scores for each sample. \*\*\**P* < 0.0005; ns, not significant (one-sided Mann-Whitney U test). (**B**) Images of the expression levels of *MIF*, *CD74*, *CD44*, and *CXCR4*, genes that constitute the MIF pathway and were expressed in ≥0.5% of all cells in each sample. (**C** and **D**) Images of spatial directions of cell-cell communication by the MIF pathway in ER_pre (C) and LR1_pre (D) samples. (**E**) Visualization of significant MIF pathway signal networks summarized by cell type in ER_pre and LR1_pre samples. (**F** and **G**) Dot plots of significant MIF pathway signal networks summarized by cell type in the ER_pre (F) and LR1_pre (G) samples. (**H** and **I**) Images of spatial directions of cell-cell communication by the PDGF pathway in ER_pre (H) and LR1_pre (I) samples. (**J**) Visualization of significant PDGF pathway signal networks summarized by cell type in ER_pre and LR1_pre samples. (**K** and **L**) Dot plots of significant PDGF pathway signal networks summarized by cell type in the ER_pre (K) and LR1_pre (L) samples.

MIF is a ligand of CD74-CD44 and CD74-CXCR4 complexes and regulates the inflammatory response^33^. To quantify ligand-receptor interactions for the MIF pathway, we calculated signal strength on the basis of the spatial proximity of ligand- and receptor-expressing cells and their expression levels. Examination of the ER_pre and LR1_pre spatial signal vectors revealed that the signals on the spatial map were directed toward the regions containing the TLSs and inflammatory cell infiltration (Fig. 6, C and D). Signal intensity and direction were further analyzed by cell type (Fig. 6, E to G).

Both ER_pre and LR1_pre samples manifested bidirectional signaling, predominantly among T cells, FDCs, and B cells. Tumor cells showed limited signals in the LR1_pre sample, whereas those in the ER_pre sample manifested signals directed toward macrophages and TLS-related cell types. Some tumor cell clusters showed activation of this pathway via an autocrine mechanism. These findings suggested that tumor cells actively stimulated macrophages and TLSs via the MIF pathway in the ER_pre sample.

Given that the proportion of fibroblasts and endothelial cells was increased after treatment for the ER samples (Fig. 2D), we next focused on the platelet-derived growth factor (PDGF) pathway, which is associated with fibroblast proliferation^34^. Signals were not localized to specific regions within the ER_pre and LR1_pre samples (Fig. 6, H and I). Bidirectional signaling was observed between fibroblasts and endothelial cells in both samples (Fig. 6, J to L). In the ER_pre sample, tumor cells, including tc_TMSB4X and tc_MMP7 clusters, showed signals directed toward stromal cells. In contrast, tumor cell clusters in the LR1_pre sample showed fewer signals directed at stromal cells. These results suggested that tumor cells interacted with stromal cells via the PDGF pathway, possibly contributing to the formation of an early-resistant TME.

## Discussion

Although various mechanisms of resistance to EGFR-TKIs in *EGFR*-mutated NSCLC have been identified, few studies have compared cases of early resistance with long-term responders^3,35,36^. We have now shown that cell proliferation in residual tumors was increased after osimertinib treatment. Genes significantly upregulated after treatment were associated with poor prognosis^37–39^, emphasizing the need to target the remaining proliferative tumor cells. Tumor cell clusters such as tc_S100A6 and tc_AQP1 were detected both before and after osimertinib treatment, suggesting that they developed resistance through activation of downstream or alternative pathways. Indeed, genes related to resistance mechanisms such as *CCND1*, *ERBB2*, and *MET* were upregulated in these clusters, with the amplification of these genes having being associated with EGFR-TKI resistance^4^. In contrast to tc_S100A6 and tc_AQP1, the tc_TMSB4X cluster was detected predominantly in the ER_pre sample and manifested low cell cycle activity, suggestive of the association of a quiescent state with EGFR-TKI resistance^40^. We found that the MIF pathway was active in the ER samples, with the tc_TMSB4X cluster manifesting activation of this pathway via an autocrine mechanism. Of note, this cluster showed significantly higher *CD74* expression compared with the other clusters. *CD74* has previously been associated with resistance to EGFR-TKIs^41^, suggesting that the tc_TMSB4X cluster may serve as a reservoir for resistant cells. Furthermore, this cluster showed increased expression of genes related to bile acid metabolism, potentially reflecting suppression of immune cell responses^42^. In contrast, the tc_SCGB3A1 cluster, which also manifested a slow cell cycle, was largely restricted to the LR1_pre sample, having disappeared after osimertinib treatment, suggestive of characteristics of low proliferation and low malignancy. The tc_MMP7 cluster was prominent in both ER_pre and ER_post samples and showed high expression of genes related to TNF-α signaling via NF-κB. Given that the activation of TNF-α signaling by osimertinib has been shown to contribute to EGFR-TKI resistance^43–45^, such signaling may constitute an early resistance mechanism. Furthermore, spatial distribution analysis revealed that the tc_MMP7 cluster was in an immunologically hot state due to a large number of plasmablasts infiltrating the surrounding area. A recent study found that CXCR1^+^ neutrophils contribute to early EGFR-TKI resistance through TNF-α signaling via NF-κB^13^. However, in our analysis, the number of neutrophils was extremely low and thus not included in the assessment. Together, our findings reveal that tumor heterogeneity contributes to EGFR-TKI resistance.

Increasing evidence implicates interactions between tumor cells and surrounding stromal or immune cells in mechanisms of EGFR-TKI resistance^9,10,46^. Cancer-associated fibroblasts have been identified as key regulators of the TME^47^. Our comparative analysis revealed an increase in the proportion of stromal cells in the ER samples after osimertinib treatment, with no marked change apparent in the SR samples and a decrease apparent in the LR samples. Of note, the proportion of fibroblasts also showed a marked increase that was accompanied by increased extracellular matrix production and upregulation of collagen genes including *COL1A1* and *COL3A1* in the ER_post sample relative to the ER_pre sample, suggestive of a potential association with early resistance^31^. *PDGFRA* and *PDGFRB*, which encode receptors for PDGF signaling, are known markers of tumor-associated fibroblasts and regulate angiogenesis^47^. PDGF signaling was active between tumor cells and stromal cells in the ER_pre sample, suggesting that tumor cells interact with stromal cells via the PDGF pathway and thereby contribute to the formation of an early-resistant TME.

We found that genes related to the inflammatory response were significantly upregulated in tumor cells of the ER-pre sample compared with SLR-pre samples. We further showed that the TME of the ER-pre sample was characterized by the presence of immature TLSs and infiltration of B cells and plasmablasts, whereas the TME of the LR1_pre sample manifested a mature TLS with reduced immune cell infiltration.

Immature TLSs have been implicated in tumor promotion, whereas mature TLSs appear to be tumor suppressive^32,48,49^. Our data suggest that the extent of TLS maturation might be a determinant of early EGFR-TKI resistance. *EGFR*-mutated NSCLC is generally characterized by low immunogenicity, which is often described as an “immune cold” state and which contributes to resistance to immunotherapy^8,9^. In contrast, an “immune hot” state, defined by TLS formation and high immune cell infiltration, has been associated with enhanced responsiveness to immunotherapy^50–54^. Of note, individuals with a short response to EGFR-TKIs have shown better responses to immunotherapy^55^, suggesting that the TME of such tumors may be characterized by an inflammatory state. Our finding that genes related to the inflammatory response were enriched in the ER_pre sample is consistent with this notion. Together, these observations indicate that an inflammatory TME can be a predictor of early resistance in *EGFR-*mutated NSCLC (Fig. 7).

**Fig. 7.**
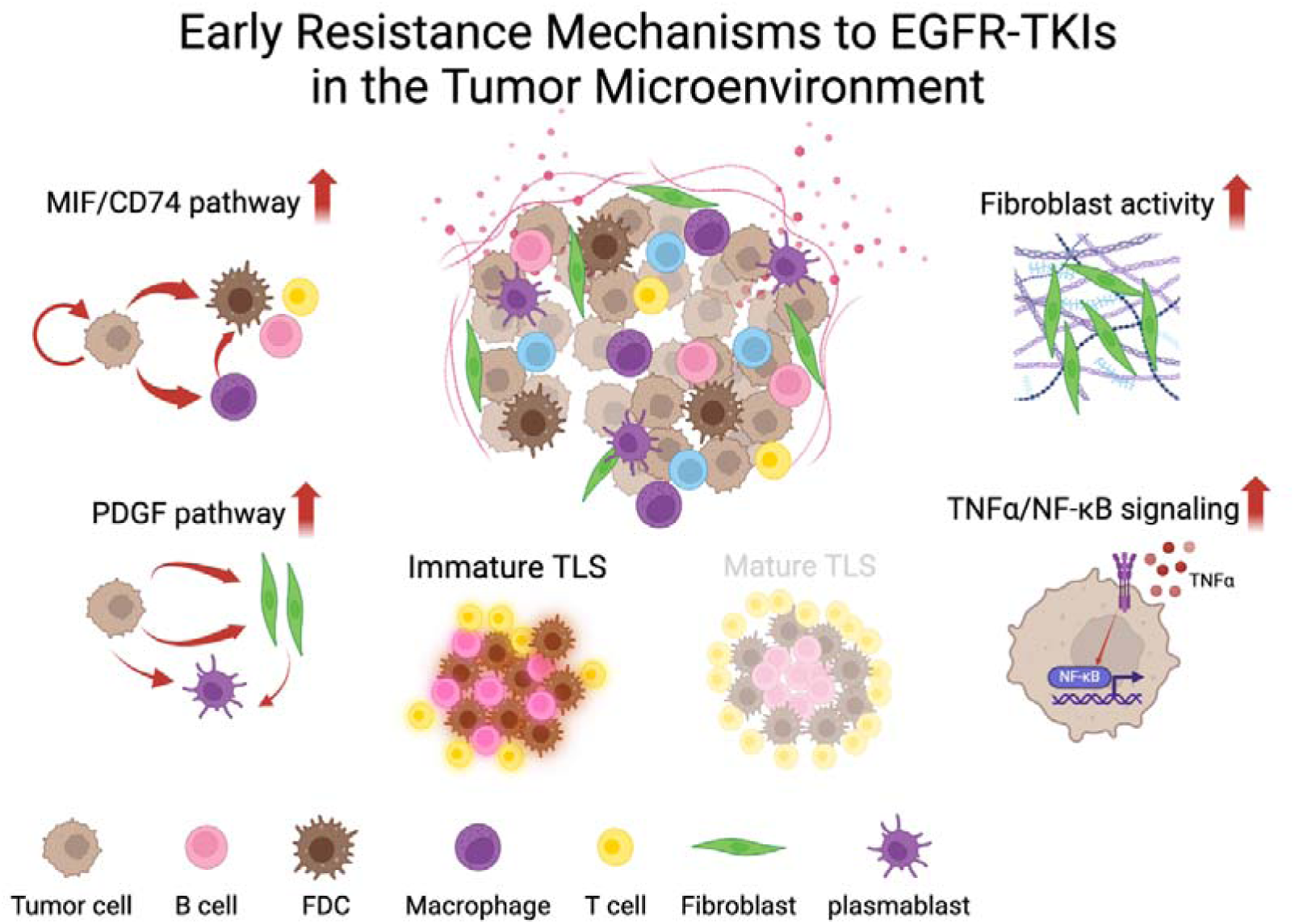
Proposed model of early resistance to EGFR-TKIs in the TME. Schematic diagram summarizing findings from the analysis of this study

Visium HD allows for high-resolution analysis, but it is difficult to distinguish between noise genes and differentially expressed genes. We addressed this issue by detecting and segmenting cell nuclei, reconstructing read count data and cell coordinate data to improve data reliability. Whereas this approach minimized false positives, some clusters still contained mixed cell types, which we subsequently excluded. As a result of these processes, comprehensive gene expression analysis and spatial analysis have become possible and the accuracy of single-cell data and spatial coordinates has improved. For further improvements in precision, multi-omics approaches are required, however.

Our study has several limitations. First, the size of the bronchoscopic lung biopsy specimens was small compared with the size of a Visium HD slide, limiting the amount of data that could be obtained. The number of tumor cells may have been insufficient to analyze the changes between before and after treatment. Second, enhancement of single-cell resolution resulted in reduced count data, potentially affecting the analysis of differentially expressed genes. Third, we cannot rule out the possibility that our findings were affected by sample variability. Future studies are warranted to collect additional clinical data for validation of our findings. In conclusion, our study suggests that an inflammatory TME and tumor heterogeneity contribute to early EGFR-TKI resistance in *EGFR*-mutated NSCLC. Of note, upregulation of resistance-related pathways was apparent in certain tumor cell clusters even before treatment onset.

## Methods

### Sample preparation

Formalin-fixed paraffin-embedded (FFPE) block specimens were collected from four *EGFR*-mutated NSCLC patients both before and after treatment with osimertinib at multiple institutions. The biopsy methods for the specimens were surgical biopsy, bronchoscopic biopsy, surgical pleural biopsy, and single-hole thoracoscopic pleural biopsy. Clinical data for the patients are provided in table S1. The patients were classified as showing early resistance (ER, *n* = 1), a standard response (SR, *n* = 1), or a long-term response (LR, *n* = 2) on the basis of the duration of PFS. The initial specimens were obtained to determine treatment. Written informed consent was obtained from all patients for specimen collection and research use. This study was performed in accordance with the Declaration of Helsinki and was approved by the Institutional Review Board of Kyushu University Hospital (approval no. 23297-01).

### Visium HD data generation

Two capture areas of a Visium HD slide (10× Genomics) were used, with four samples being assigned to each capture area. A total of eight samples consisting of paired pre-and posttreatment specimens from four patients (ER_pre, ER_post, SR_pre, SR_post, LR1_pre, LR1_post, LR2_pre, LR2_post) was analyzed, with ER and SR samples being assigned to one capture area and LR1 and LR2 samples to the other. RNA was extracted from the FFPE tissue sections for RNA quality assessment, and the DV200 (percentage of RNA fragments of >200 nucleotides) value was measured with the Agilent 4200 TapeStation instrument. The DV200 value for all samples was ≥30%. FFPE tissue sections were then depleted of paraffin and subjected to H&E staining according to the Visium HD FFPE Tissue Preparation protocol (Visium HD FFPE Tissue Preparation Handbook, CG000684 Rev A). The tissue sections were imaged with an Evident VS200 microscope in accordance with the 10× Genomics protocol (Visium HD Spatial Applications Image Guidelines CG000688 Rev A). Gene expression libraries were prepared from the H&E-stained sections according to the 10× Genomics protocol (Visium HD Spatial Gene Expression Reagent Kits User Guide CG000685 Rev A). The quality of the prepared libraries was evaluated with the Agilent 4200 TapeStation instrument, and all libraries were confirmed to have a sufficient concentration. The Visium libraries were sequenced with the Illumina NovaSeq X Plus platform with a paired-end configuration (read 1, 150 bp; read 2, 150 bp). Sequencing data (FASTQ files) were processed and analyzed with Space Ranger (version 3.0.0) software for assessment of data quality and generation of spatial gene expression profiles.

### Analysis environment

Analyses were conducted with Python (version 3.10) on either Ubuntu (22.04.5 LTS) with JupyterLab (4.2.5) or Windows (Windows 11 Home Edition) with PyCharm (Community Edition 2024.1.1). The primary Python libraries and their versions included AnnData (0.10.8), AnnData2ri (1.3.2), Commot (0.0.3), GeoPandas (1.0.1), GSEApy (1.1.3), HarmonyPy (0.0.10), inferCNVpy (0.5.0), Matplotlib (3.8.4), NumPy (1.26.4), Pandas (2.2.2), Rpy2 (3.5.16), Scanpy (1.10.3), SciPy (1.11.4), Seaborn (0.13.2), Squidpy (1.6.1) and StarDist (0.9.1). For some analyses, R Script was executed in Python. The primary R packages and their versions used in this study were tradeSeq (1.8.0), edgeR (3.36.0), and clusterExperiment (2.14.0). Whereas the libraries were generally applied as described in their documentation, modifications were made to Commot library to improve visual clarity in plots.

### Detection and segmentation of nuclei

Although Visium HD allows for the use of 8-µm square spots as single cells, these spots do not accurately reflect true cellular coordinates. For preservation of cellular coordinates as much as possible, StarDist was applied for detection and segmentation of nuclei^19^. Specifically, Python code was developed on the basis of the Analysis Guide provided by 10× Genomics (https://www.10xgenomics.com/jp/analysis-guides/segmentation-visium-hd). Running of this code generated coordinate data for individual nuclei represented as polygons. Count data for each 2-µm spot within the polygon area were relabeled as corresponding to a single cell. In addition, the centroid coordinates of each polygon were added to the count data, together with metadata content, thereby generating a data structure compatible with libraries commonly used in spatial transcriptomics analysis.

### Sample labeling on spatial coordinates

In typical spatial transcriptomics analysis, all samples within a single capture area are assumed to be the same. However, in the present study, four different samples were present within each capture area. Given that this situation is not typically supported by existing tools, it was necessary to label samples manually on the basis of spatial coordinates. With the use of the spatial coordinate data within the data structure, samples were divided into eight groups by referencing images and assigning coordinates manually. Given that there was an overlapping area between LR1_pre and LR1_post specimens, a blank area was generated to avoid sample contamination.

### Major clustering

The newly generated data from nuclei segmentation was analyzed with a pipeline similar to that for single-cell RNA-seq analysis with the Scanpy package. Cells with <20 counts or containing >10% mitochondrial genes as well as genes with <20 total counts were excluded from the analysis. Cells with a cross-sectional area of ≥900 µm^2^ on the image were excluded given the possibility that they might actually have been composed of multiple cells. After filtering, the data were calculated for highly variable genes (HVGs) and normalized with Pearson residuals, which adjusts for library size and overdispersion. Principal component analysis (PCA) for the top 4000 HVGs allowed calculation of principal components up to the top 35. Harmony integration was applied with the use of the HarmonyPy library through Scanpy with the sample labels in order to account for batch effects across samples. The nearest-neighbors distance matrix and the neighborhood graph of the observations were then calculated with a local neighborhood size of 100 and 35 principal components. An UMAP embedding was generated with a minimum distance parameter set to 0.5. Leiden clustering was performed at a resolution of 0.4 and identified 18 clusters. However, given the high noise level in cells, it was possible that the differences between cells and noise would not be distinguishable. Instead, we analyzed the expression of HVGs in each cluster and compared the expression levels of known cell markers to classify the clusters into three major cell types: epithelial, stromal, and immune (table S3).

### Subcluster analysis

Subclustering was performed for further exploration of the epithelial, stromal, and immune cell populations identified in the initial clustering. For each cell type cluster, the analysis was started again from the raw count data. The data were filtered to the same threshold as for the major clustering. HVGs were calculated within some subclusters and PCA was performed, followed by the construction of neighborhood graphs, UMAP embedding, and clustering. The distinction between tumor cells and normal cells was evaluated with inferCNVpy, which is based on inferCNV, in addition to the pathological images. Normal airway epithelial cells obtained from the clustering results were set as nonmalignant cells and analyzed. Furthermore, additional clustering was performed for tumor cells. For immune cells, three cell types—lymphocytes, plasmablasts, and macrophages—were identified, and each type was subsequently analyzed with individual clustering steps. Detailed parameters for each clustering step and cell annotation markers are provided in tables S3 and S4.

### DGE Analysis

DGE analysis was performed with generated pseudo–bulk samples and the edgeR package^56,57^. Single-cell RNA-seq data were aggregated at the sample level for each cell type of interest. Samples with <30 cells of the target cell type were excluded. For comparisons between ER and SLR groups as well pre-versus posttreatment comparisons of the tc_AZGP1 cluster, two random replicates were generated for each group because the edgeR library requires biological replicates for statistical modeling. For other comparisons, pseudo–bulk aggregation was performed without generation of additional replicates. After pseudo–bulk aggregation, DGE analysis was performed with the edgeR library^57^. Low-expression genes were first filtered out, retaining only those with sufficient counts across experimental groups to improve statistical power. Library size normalization was performed with the trimmed mean of M-values method, which adjusts for sequencing depth and compositional biases across samples. A design matrix was constructed to account for sample conditions and randomly generated replicates, allowing the estimation of both biological variation and group-specific effects. The quasi–likelihood variance estimation method was adopted to generate a gene list for DGE. Adjusted *P* values were calculated with the Benjamini-Hochberg method to control the false discovery rate (FDR). Results for each cell type were integrated back into the annotated data object to facilitate further interpretation. The aggregated gene expression values were subsequently used for downstream analyses, including the construction of volcano plots and GSEA.

### Kaplan-Meier Plotter database

We used data from lung cancer patients in the Kaplan-Meier Plotter database (http://kmplot.com/analysis) to evaluate differentially expressed genes in tumor cells compared before and after treatment^58^. Patients in the database were divided into high-and low-expression groups on the basis of median expression values, and overall survival curves were plotted with no follow-up restrictions.

### Integration of cell type and spatial coordinate data

The data set for this analysis was generated by integration of single-cell analysis for cell type identification with spatial coordinate data derived from StarDist. The StarDist-based coordinate data, which represent new spot coordinates, were transformed on the basis of the ratio of the actual distance to the pixel coordinates, as described in the metadata. This integration resulted in a unified data structure compatible with analysis tools such as Scanpy and Squidpy.

### Neighborhood enrichment analysis

Neighborhood enrichment analysis was performed on the spatial transcriptomics data to assess the enrichment of different cell clusters within their spatial neighborhoods. For each of the eight sample data sets, the spatial neighbors function within Squidpy was used to calculate the spatial neighbors of each spot. The neighborhood enrichment function was then applied to calculate the z-scores with predefined cell cluster labels.

For standardization of the enrichment scores across all samples, the minimum and maximum values were first calculated across all data sets. Each z-score matrix was then normalized by subtracting the global mean and dividing by the standard deviation, ensuring that enrichment scores were comparable across samples. This normalization allowed for the unified analysis of neighborhood enrichment across different experimental conditions.

### Spatial distribution and neighborhood analysis of tumor cells

The spatial distribution and neighborhood relations of tumor cells were analyzed by integration of single-cell cluster information with spatial coordinate data. For each of the eight sample data sets, the spatial positions of tumor cells were identified. The composition of surrounding cells was quantified by calculation of the pairwise distances between each tumor cell and all other cell types within the tissue. For analysis of the spatial distribution, concentric annular regions extending up to 1000 µm from each tumor cell were defined, with each region increasing in 20-µm increments. These regions served as distance bins, allowing categorization of surrounding cells on the basis of their proximity to tumor cells. The proportion of each cell type within these bins was then calculated to assess the spatial distribution relative to tumor clusters. This method allowed a quantitative evaluation of neighborhood enrichment patterns, providing insight into the TME and its cellular composition.

### Cell-cell communication analysis

Cell-cell interactions were analyzed with COMMOT, a tool that allows comprehensive modeling of both direct and indirect interactions between different cell types while incorporating the spatial organization of cells within tissues^59^. Smooth muscle cells, normal airway epithelial cells, and blood cells were excluded from this analysis to allow focus on the TME. In addition, genes expressed in ≥0.5% of all cells were first filtered. The gene list was further refined by extraction of known ligand-receptor pairs from the CellChat database. Only ligand-receptor pairs co-occurring within a distance of 500 µm were considered for subsequent analysis. For the identified ligand-receptor pairs and their associated pathways, signal strength and vectors were calculated on the basis of the distance between sending and receiving cells as well as gene expression intensity.

### Gene expression analysis and statistics

Gene expression scores were calculated by a marker gene–based scoring method implemented in Scanpy. The MIF score was derived from the MIF pathway–related genes *MIF*, *CD74*, *CD44*, and *CXCR4*. Background correction was performed with control gene sets of 5000 randomly selected genes to ensure robust scoring. Pairwise comparisons of the MIF score and of gene expression in tumor cells between samples were performed with the unilateral Mann-Whitney U test. Visualization was conducted with Seaborn for violin and strip plots annotated with the *P* value. The significance level for GSEA after DGE analysis was set as a *P* value of <0.05 and FDR of 0.25.

## Data availability

All data associated with this study are in the paper or the Supplementary Materials. The Visium HD data set for this study is available in the GEO repository under accession number GSE288758.

## Acknowledgments

We thank CyberomiX (Kyoto, Japan) for assistance with spatial transcriptomics analysis via Visium HD platform.

## Author contributions

D.S., K.T., and I.O. made substantial contributions to the conception of the work. Y.K., K.N., N.N., M.K., and K.A. provided samples and clinical information for this work. S.N., D.S., K.T, M.H., R.I., K.O., Y.Y., E.I., and I.O. made significant contributions to data interpretation. S.N. and D.S. analyzed the data and drafted the original manuscript. S.N. and D.S. reviewed and edited the manuscript. Y.O. and I.O. supervised the study.

