## Supplementary information for "Spatial dynamics of the tumor microenvironment linked to emerging resistance in *EGFR*-mutated lung cancer"

**Supplementary Data**

**Supplementary Figures:**

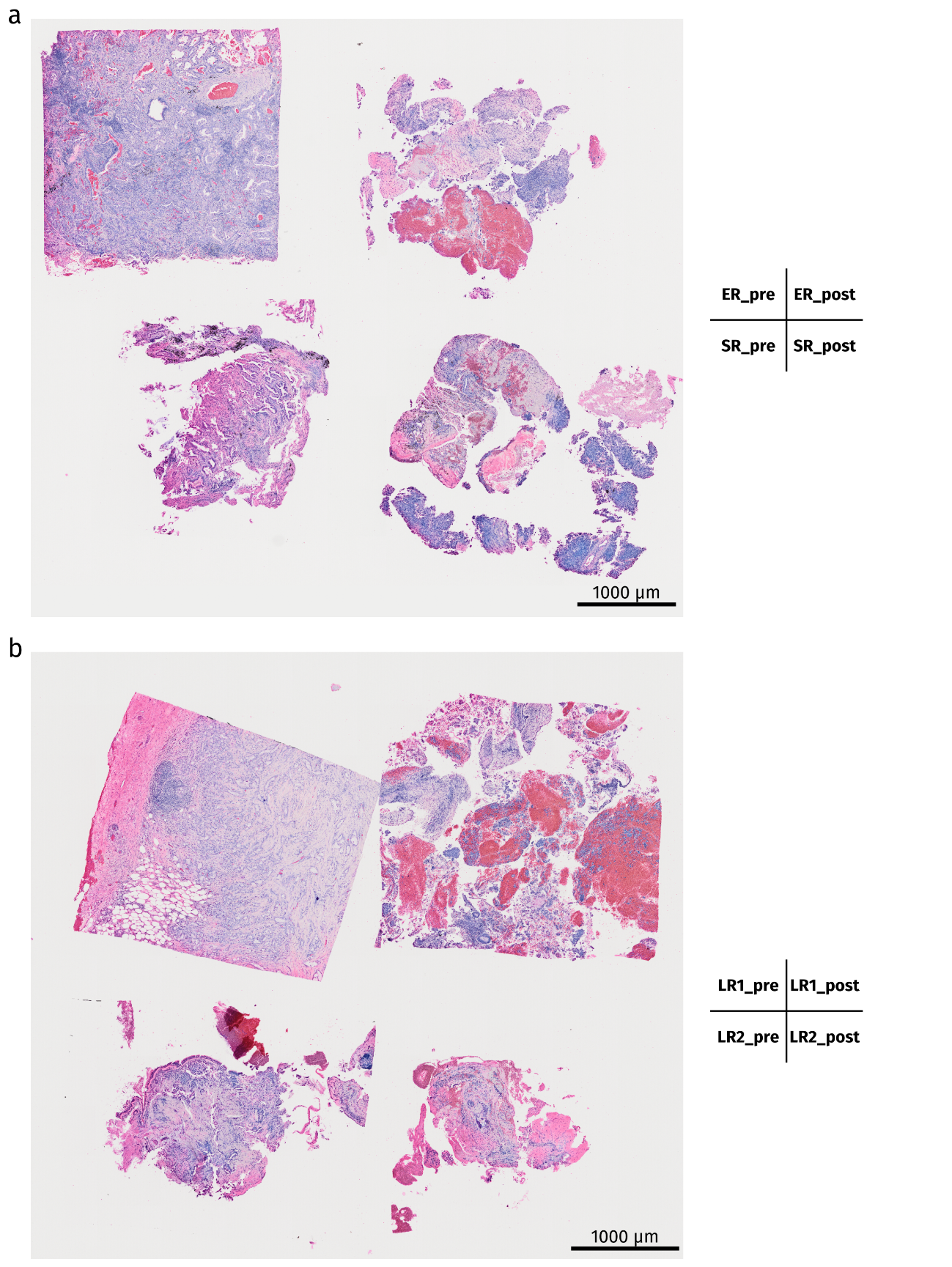

**Supplementary Fig. 1 | H&E staining of samples on Visium HD slides. a**, Pre- and posttreatment samples from the patients classified as ER and SR on Slide 1. **b**, Pre- and posttreatment samples from the patients classified as LR1 and LR2 on Slide 2.

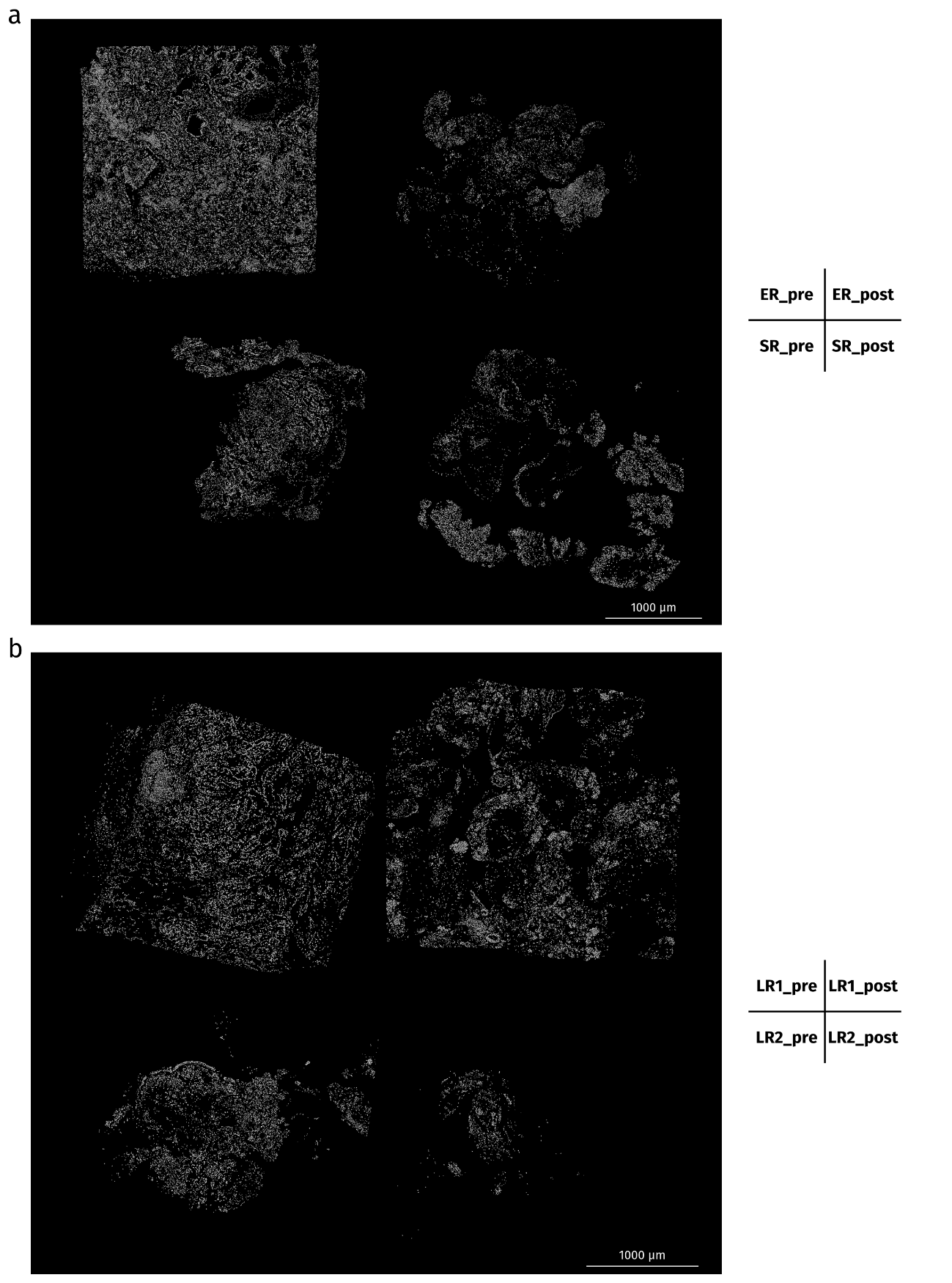

**Supplementary Fig. 2 | Cell nucleus images of Slides 1 and 2 obtained with StarDist.** Each cell was plotted on the basis of the polygon coordinate data obtained by StarDist processing. A total of 141,324 cell nuclei was identified in Slide 1 (**a**) and 99,658 cell nuclei in Slide 2 (**b**).

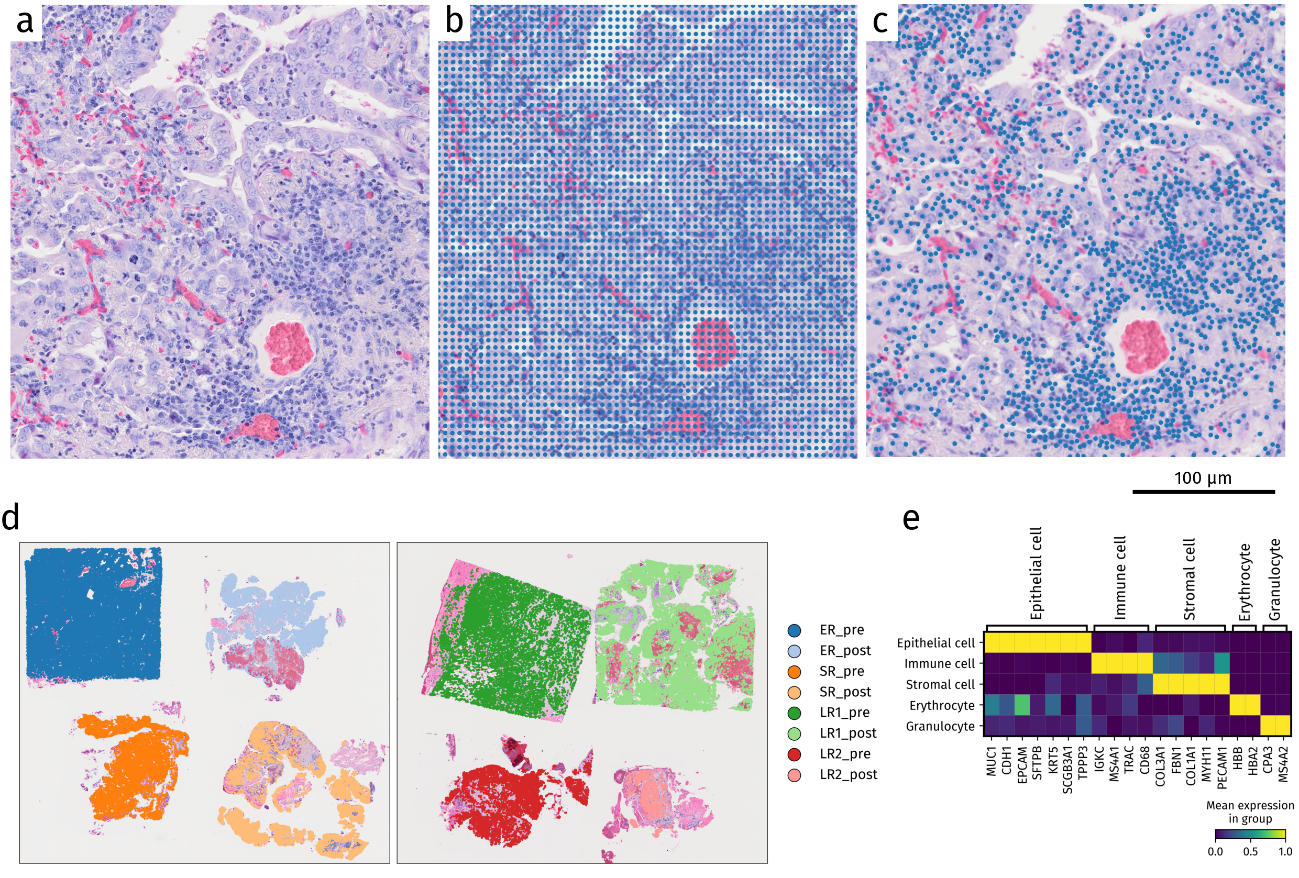

**Supplementary Fig. 3 | Sample identification and clustering based on the integration of new coordinates and new cell data. a**, H&E-stained pathological image of an area of ER_pre. **b**, Eight-micrometer spot original coordinates plotted on the H&E-stained image. The spots were independent of cell location and were present even in areas where no cells were located. **c**, New spots plotted at the centroids of the polygons derived by StarDist. These spots are consistent with the actual positions of the cells, allowing more rigorous spatial analysis. **d**, Identification of samples in the data based on coordinates in the spot data and tissue images. **e**, Matrix plot of cell markers used to identify cell types and each major cluster.

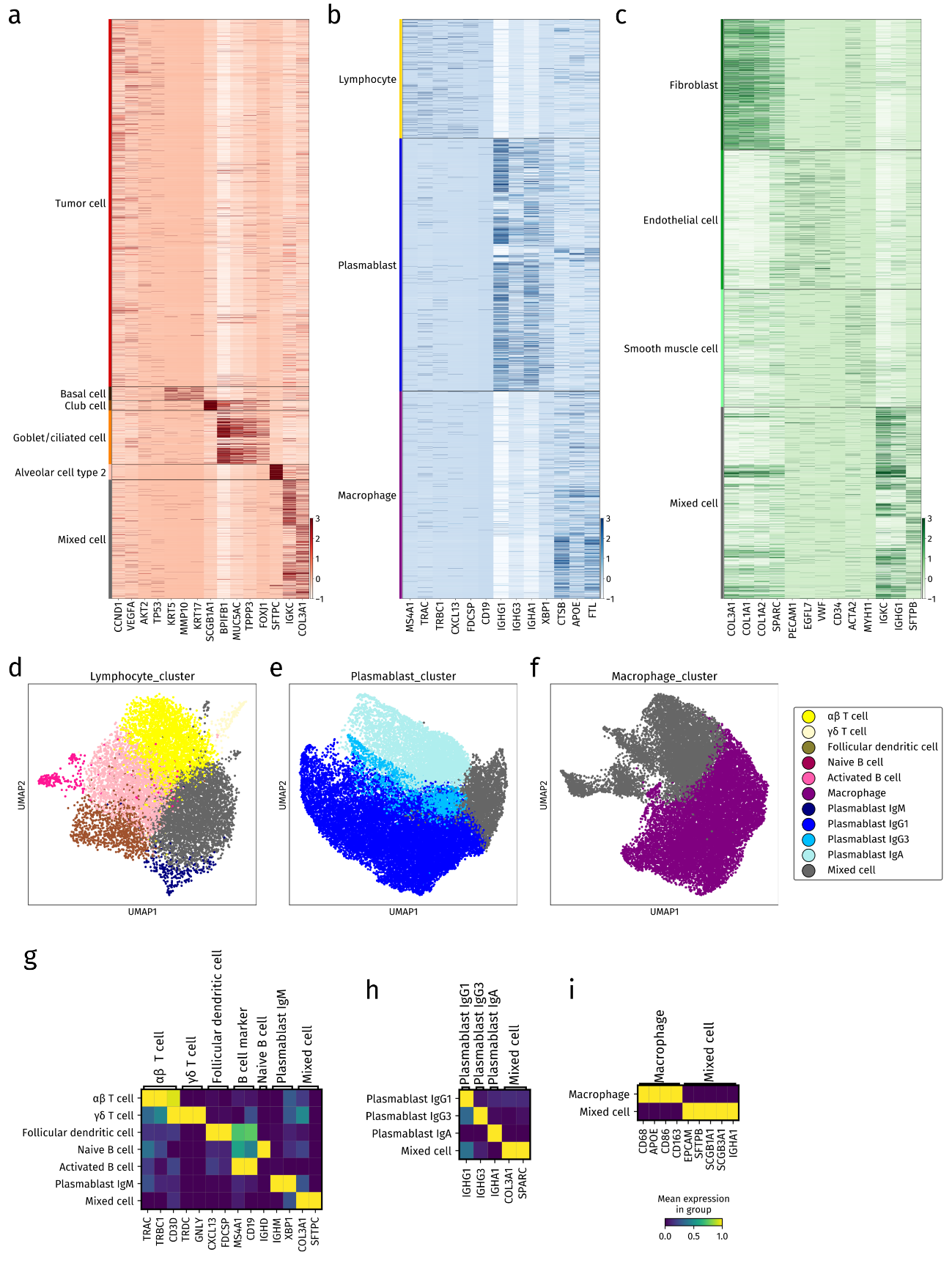

**Supplementary Fig. 4 | Subcluster analysis of the three major lineages and identification of cell types. a–c**, Heat maps showing the expression of marker genes and cell types identified for epithelial cells (**a**), immune cells (**b**), and stromal cells (**c**). Goblet cells and ciliated cells could not be distinguished because many cells had characteristics of both. **d–f**, UMAP of cell types in lymphocyte (**d**), plasmablast (**e**), and macrophage (**f**) clusters. **g–i**, Matrix plots of identified cell types and marker genes for lymphocyte (**g**), plasmablast (**h**), and macrophage (**i**) clusters.

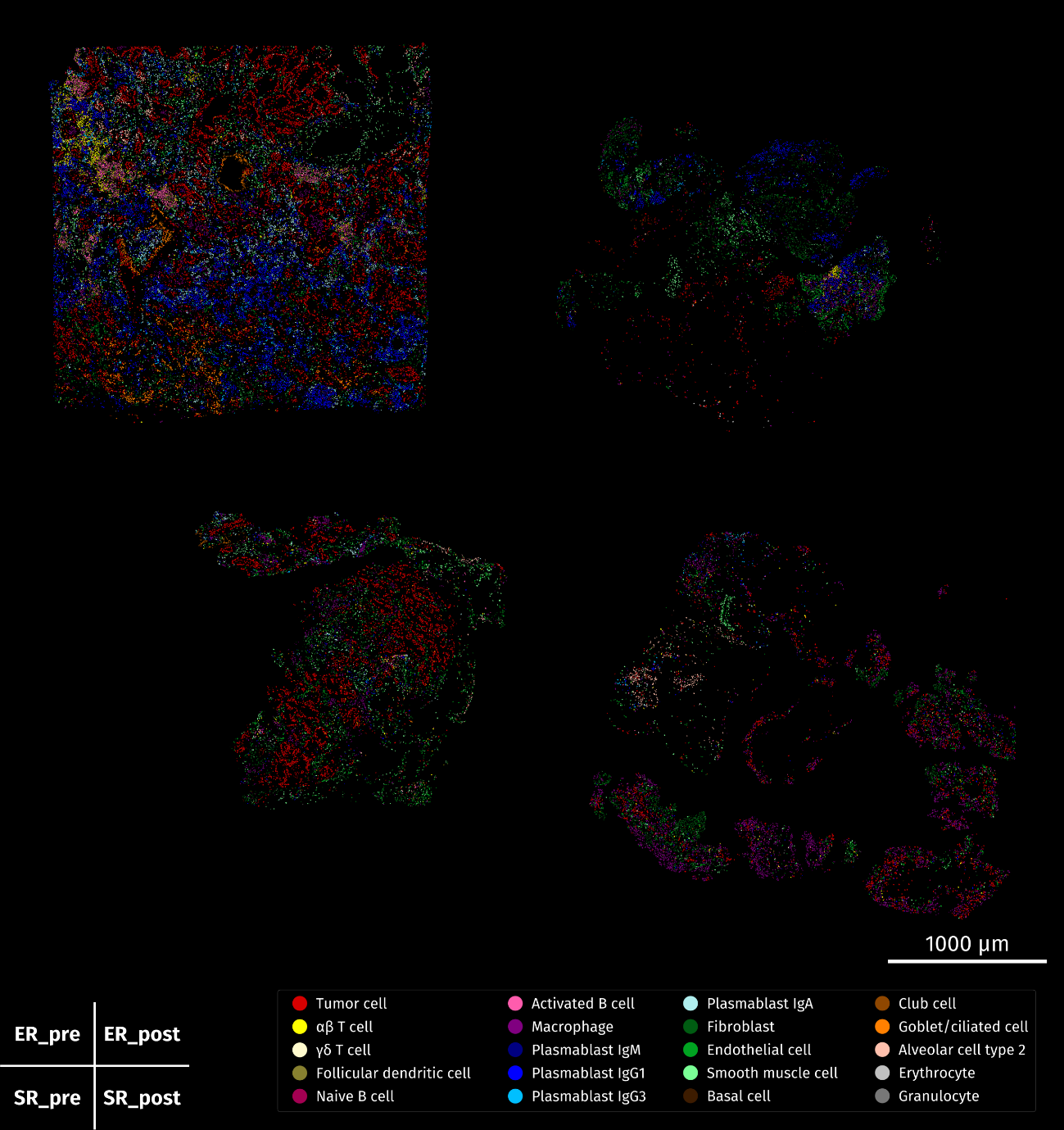

**Supplementary Fig. 5 | Plot image of Slide 1 based on polygon coordinates.** The cell types obtained by clustering were integrated into the polygon coordinates derived by StarDist processing, and Slide 1 area was plotted.

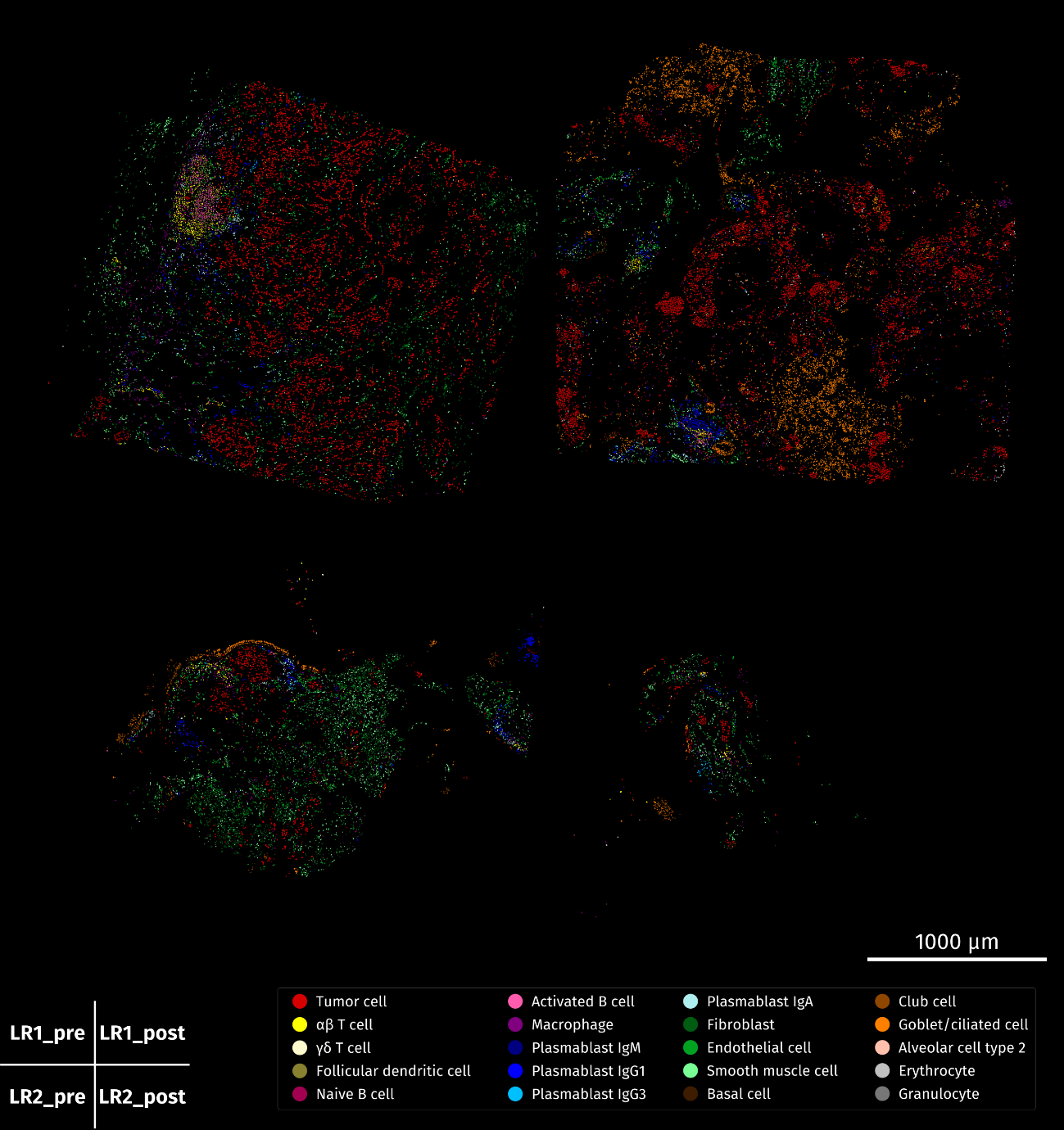

**Supplementary Fig. 6 | Plot image of Slide 2 based on polygon coordinates.** The cell types obtained by clustering were integrated into the polygon coordinates derived by StarDist processing, and Slide 2 area was plotted.

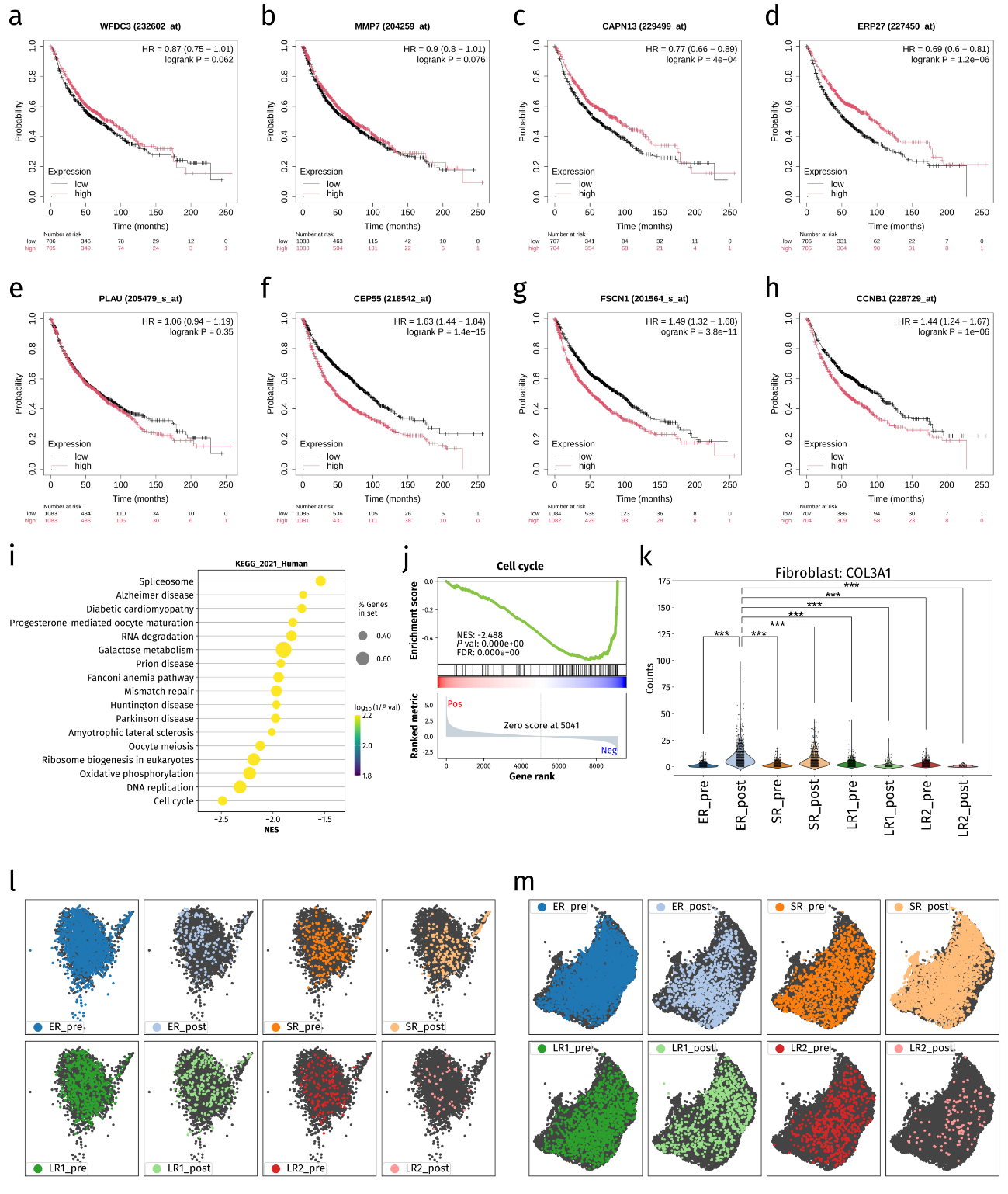

**Supplementary Fig. 7 | Comparative analysis of samples obtained before and after treatment. a–h**, Prognostic estimation for NSCLC patients in Kaplan-Meier Plotter according to the expression level of genes whose expression was significantly increased in All_pre (**a–d**) and All_post (**e**–**h**) samples in the DGE analysis of tumor cells for All_pre vs All_post. The y-axis represents overall survival probability, and the x-axis shows time in months. *P* values were calculated by log-rank test. HR, hazard ratio (95% confidence interval shown in parentheses). **i**, Dot plot for GSEA with KEGG (Kyoto Encyclopedia of Genes and Genomes) gene sets based on DGE analysis of tumor cells for All_pre vs All_post. **j**, GSEA plot for a gene set with a high negative value of NES for tumor cells of All_pre vs. All_post. **k**, Violin plot and strip plot showing raw counts for the *COL3A1* gene in fibroblasts for each sample. ****P* < 0.0005 (one-sided Mann-Whitney U test). **l**, Distribution of each sample shown on the UMAP plot for the T cell part extracted from the lymphocyte cluster. **m**, Distribution of each sample shown on the UMAP plot of the macrophage cluster.

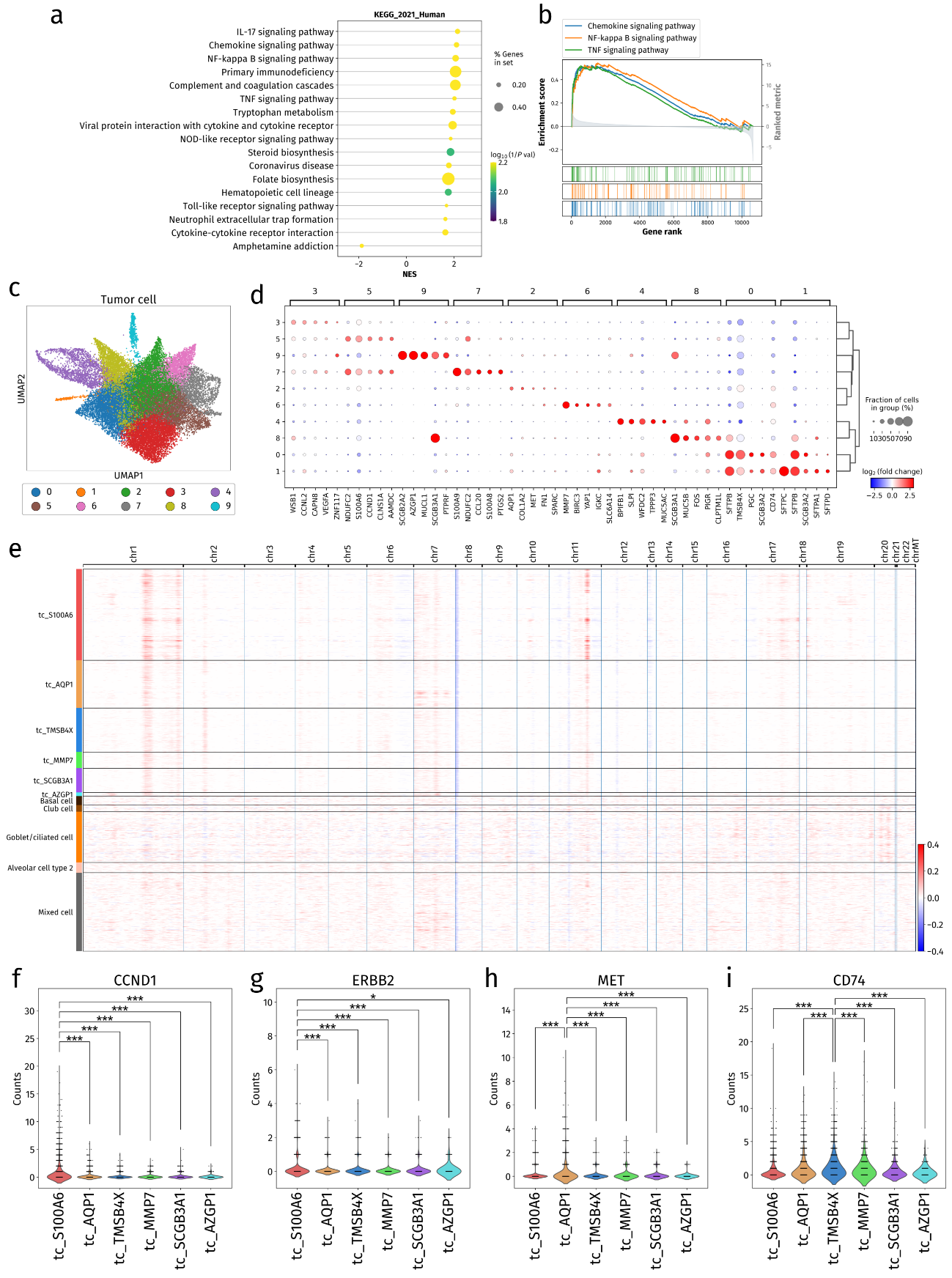

**Supplementary Fig. 8 | Analysis exploring mechanisms underlying tumor heterogeneity and early resistance. a**, Dot plot for GSEA with KEGG gene sets based on DGE analysis of tumor cell clusters of ER_pre vs SLR_pre. **b**, GSEA plots for gene sets with high positive values of NES for ER_pre vs SLR_pre. **c**, UMAP plot showing tumor cells reclustered and labeled for Leiden clustering. **d**, Dot plot for the top five differentially expressed genes in each Leiden clustering. **e**, Heat map of epithelial cells analyzed by inferCNVpy. Basal cells, club cells, goblet/ciliated cells, and alveolar cells type 2 were set as nonmalignant cells. Concordant CNV changes were predicted in tumor cells compared with normal cells. **f**–**i**, Violin plots and strip plots for raw counts of *CCND1* (**f**), *ERBB2* (**g**), *MET* (**h**), and CD74 (**i**) for each tumor cell cluster. **P* < 0.05, ****P* < 0.0005 (one-sided Mann-Whitney U test).

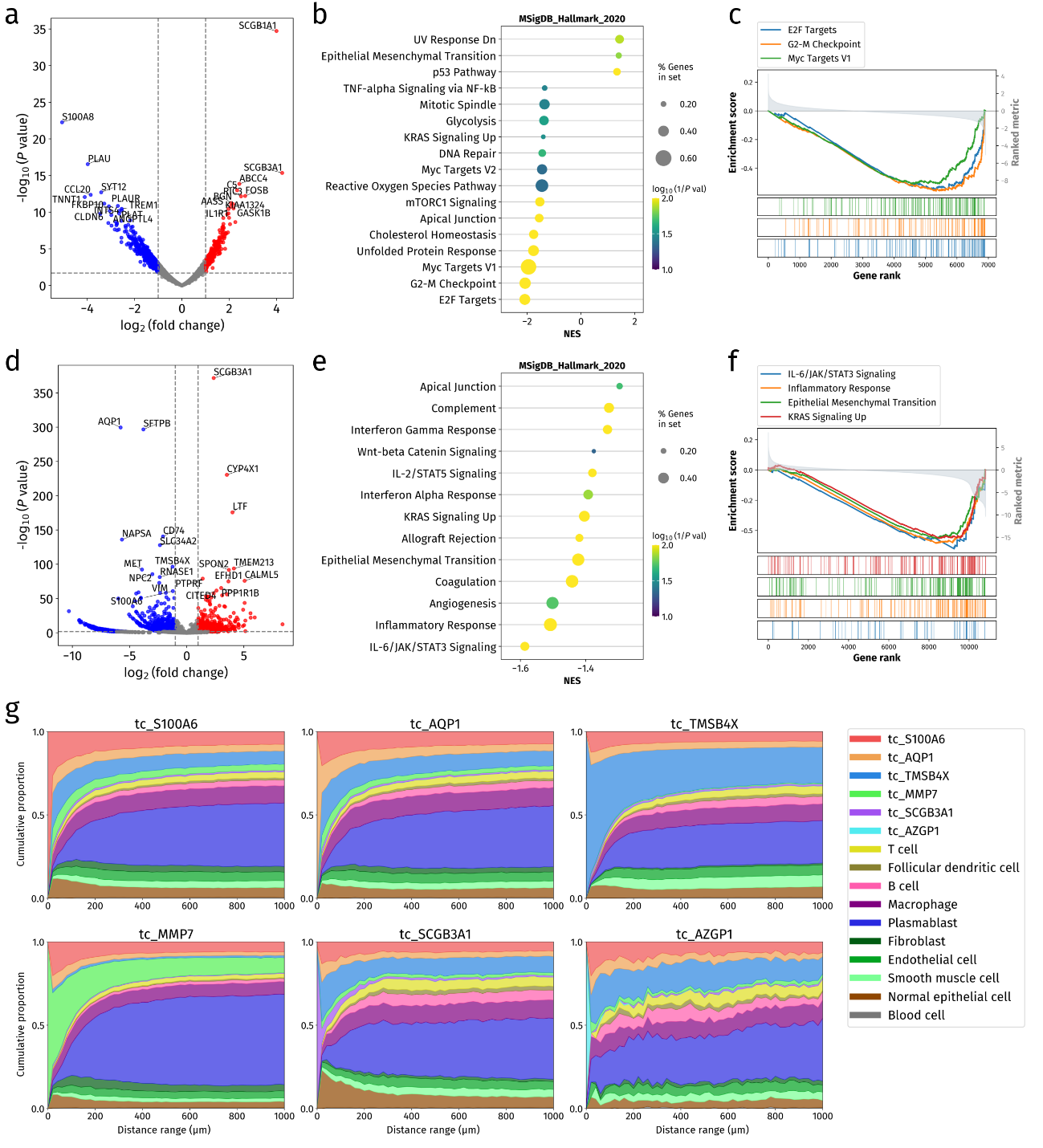

**Supplementary Fig. 9 | Analysis of individual tumor cell clusters and cluster composition. a**, Volcano plot of DGE analysis results for the tc_SCGB3A1 cluster versus the corresponding other tumor cell clusters (tc_rest). **b**, Dot plot for GSEA with Hallmark gene sets based on DGE analysis of tc_SCGB3A1 vs tc_rest. **c**, GSEA plots for gene sets with high negativevalues of NES for tc_SCGB3A1 vs tc_rest. **d**, Volcano plot of DGE analysis results for tc_AZGP1 vs tc_rest. **e**, Dot plot for GSEA with Hallmark gene sets based on DGE analysis of tc_AZGP1 vs tc_rest. **f**, GSEA plots for gene sets with high negative values of NES for tc_AZGP1 vs tc_rest. **g**, Stacked line graph diagrams depicting the quantified spatial distribution of cell types around tumor cells in the ER_pre sample. For each tumor cell cluster, the proportion of cell types within concentric ring regions extending up to 1000 μm from each tumor cell is shown.

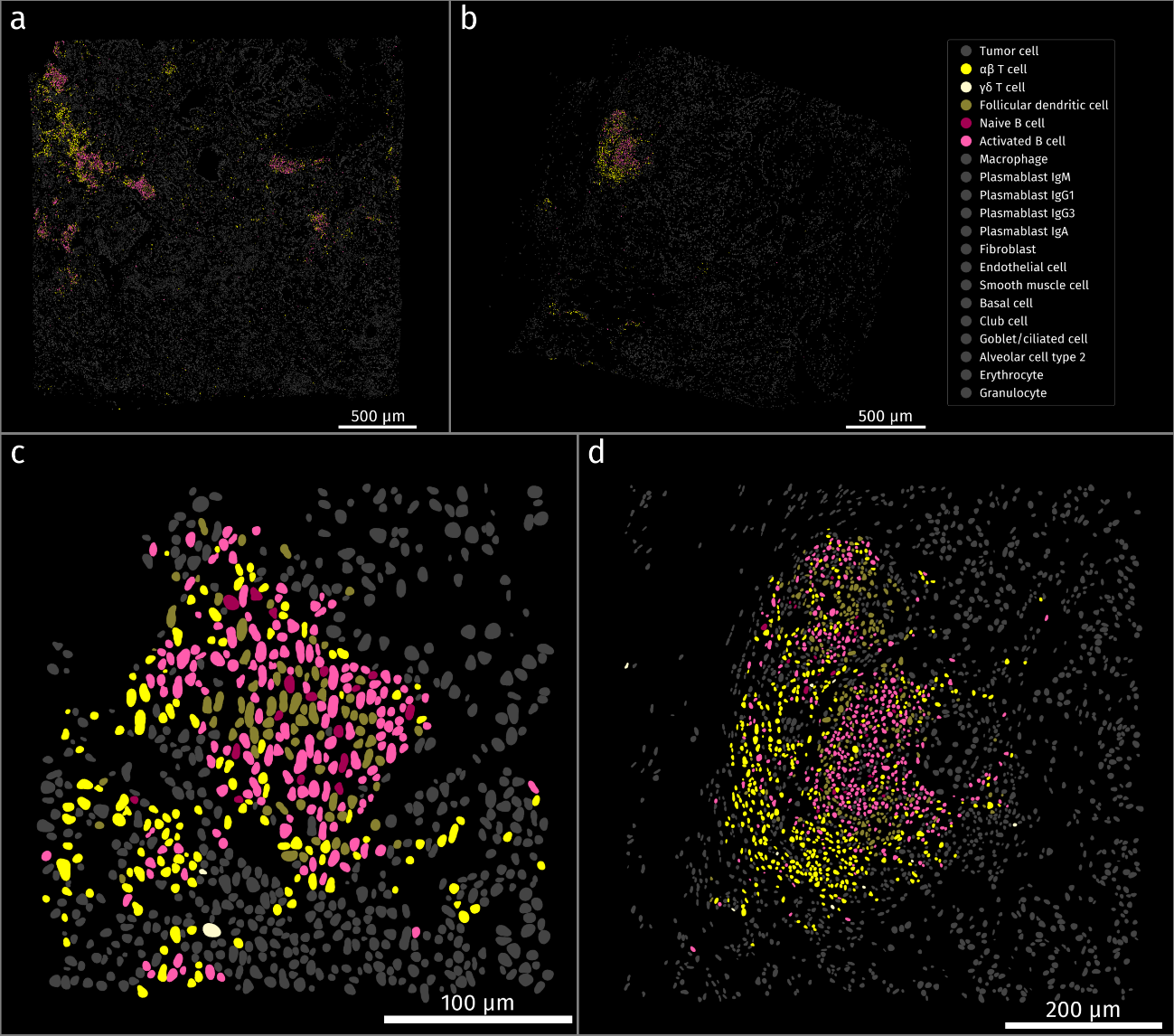

**Supplementary Fig. 10 |** **Difference in TLS maturity between samples. a**,**b**, Images of the entire ER_pre (**a**) and LR1_pre (**b**) samples showing only the coloring of T cells, B cells, and follicular dendritic cells (FDCs). **c**,**d**, Enlarged images of the TLSs in the ER_pre sample (**c**) and the LR1_pre sample (**d**) showing only the coloring of T cells, B cells, and FDCs.

**Supplementary Tables:**

**Supplementary Table 1 | Patient background for specimens collected for analysis.**

| Characteristic | ER | SR | LR1 | LR2 |
| --- | --- | --- | --- | --- |
| Age (years) | 75 | 63 | 75 | 75 |
| Sex | Female | Male | Male | Male |
| Smoking history | Never | Never | Former | Former |
| Histological type | Adenocarcinoma | Adenocarcinoma | Adenocarcinoma | Adenocarcinoma |
| *EGFR* mutation | Exon-19 deletion | L858R | Exon-19 deletion | L858R |
| PD-L1 TPS (%) | 1–24 | 1–24 | 5 | <1 |
| Metastasis | OSS | OSS, BRA, HEP  LYM, PLE | PLE | PLE, HEP |
| Biopsy pretreatment | Surgical biopsy | TBLB | Surgical pleural biopsy | TBLB |
| Biopsy posttreatment | Pleural biopsy | TBLB | TBLB | TBLB |
| Osimertinib treatment period (months) | 6 | 16 | 36 | 24 |

PD-L1, programmed cell death–ligand 1; TPS, tumor proportion score; OSS, osseous (bone) metastasis; BRA, brain metastasis; HEP, hepatic metastasis; LYM, lymph node metastasis; PLE, pleural metastasis; TBLB, transbronchial lung biopsy.

**Supplementary Table 2 | Marker genes for cell type identification.** Detection of cell types of each cluster was based on these marker genes and differentially expressed genes.

| Cluster | Subcluster | Marker genes |
| --- | --- | --- |
| Major cluster | Epithelial cell | *MUC1, CDH1, EPCAM, SFTPB, KRT5, SCGB3A1, TPPP3* |
|  | Immune cell | *IGKC, MS4A1, TRAC, CD68* |
|  | Stromal cell | *COL3A1, FBN1, COL1A1, MYH11, PECAM1* |
|  | Erythrocyte | *HBB, HBA2* |
|  | Granulocyte | *CPA3, MS4A2* |
| Epithelial cell | Tumor cell | *CCND1, VEGFA, AKT2, TP53* |
|  | Basal cell | *KRT5, MMP10, KRT17* |
|  | Club cell | *SCGB1A1* |
|  | Goblet/ciliated cell | *BPIFB1, MUC5AC, TPPP3, FOXJ1* |
|  | Alveolar cell type 2 | *SFTPC* |
|  | Mixed cell | *IGKC, COL3A1* |
| Immune cell | Lymphocyte | *MS4A1, CD19, TRAC, TRBC1, CXCL13, FDCSP* |
|  | Plasmablast | *IGHG1, IGHG3, IGHA1, XBP1* |
|  | Macrophage | *CTSB, APOE, FTL* |
| Stromal cell | Fibroblast | *COL3A1, COL1A1, COL1A2, SPARC* |
|  | Endothelial cell | *PECAM1, EGFL7, VWF, CD34* |
|  | Smooth muscle cell | *ACTA2, MYH11* |
|  | Mixed cell | *IGKC, IGHG1, SFTPB* |
| Lymphocyte | αβ T cell | *TRAC, TRBC1, CD3D* |
|  | γδ T cell | *TRDC, GNLY, CD3D* |
|  | Follicular dendritic cell | *CXCL13, FDCSP* |
|  | Naive B cell | *MS4A1, CD19, IGHD* |
|  | Activated B cell | *MS4A1, CD19* |
|  | Plasmablast IgM | *IGHM, XBP1* |
|  | Mixed cell | *COL3A1, SFTPC* |
| Plasmablast | Plasmablast IgG1 | *IGHG1* |
|  | Plasmablast IgG3 | *IGHG3* |
|  | Plasmablast IgA | *IGHA1* |
|  | Mixed cell | *COL3A1, SPARC* |
| Macrophage | Macrophage | *CD68, APOE, CD86, CD163* |
|  | Mixed cell | *EPCAM, SFTPB, SCGB1A1, SCGB3A1, IGHA1* |

**Supplementary Table 3 | Parameters of single-cell RNA sequence for each cluster.**

| Cluster | HVGs | PCs | n_neighbor | n_pcs | resolution | min_dist |
| --- | --- | --- | --- | --- | --- | --- |
| Epithelial cell | Top 1000 genes | 20 | 100 | 20 | 0.6 | 0.4 |
| Immune cell | Not used | 30 | 30 | 10 | 0.4 | 0.4 |
| Stromal cell | Top 2000 genes | 30 | 80 | 6 | 0.3 | 0.3 |
| Tumor cell | Not used | 50 | 100 | 10 | 0.5 | 0.5 |
| Lymphocyte | * | * | 150 | 30 | 0.7 | 0.5 |
| Plasmablast | * | * | 20 | 8 | 0.3 | 0.5 |
| Macrophage | * | * | 80 | 10 | 0.3 | 0.5 |

*Lymphocytes, plasmablasts, and macrophages were reclustered with calculated principal components (PCs) in immune cell clustering. HVGs, highly variable genes; n_neighbor, local neighborhood size; n_pcs, number of PCs; resolution, Leiden clustering parameter; min_dist, UMAP embedding parameter.

**Supplementary Table 4 | Ligands, receptors, and pathways extracted on the basis of the number of gene-expressing cells.** Ligand-receptor pairs and the corresponding pathways for signal transduction expressed in ≥0.5% of all cells in the ER_pre sample are listed.

| Ligand | Receptor | Pathway |
| --- | --- | --- |
| GDF15 | TGFBR2 | GDF |
| PDGFA | PDGFRA | PDGF |
| PDGFA | PDGFRB | PDGF |
| CXCL12 | CXCR4 | CXCL |
| MIF | CD74_CXCR4 | MIF |
| MIF | CD74_CD44 | MIF |
| CSF1 | CSF1R | CSF |
| TNFSF12 | TNFRSF12A | TWEAK |
| MDK | SDC1 | MK |
| MDK | SDC4 | MK |
| MDK | ITGA4_ITGB1 | MK |
| MDK | LRP1 | MK |
| C3 | ITGAX_ITGB2 | COMPLEMENT |
| GAS6 | AXL | GAS |
| GRN | SORT1 | GRN |
| LGALS9 | CD44 | GALECTIN |
